# Distributed burst activity in the thalamocortical system encodes reward contingencies during learning

**DOI:** 10.1101/2025.03.11.642368

**Authors:** Filippo Heimburg, Josephine Timm, Nadin Mari Saluti, Alexander Groh

## Abstract

Neuronal bursts are distinct high-frequency firing patterns that are present ubiquitously throughout mammalian brain circuits. Although bursts are considered part of a universal neural code, the information they convey has long been a subject of debate. In this study, we investigated neuronal activity in simultaneously recorded regions of the thalamocortical system in freely moving mice as they learned stimulus-outcome associations in a go/no-go task. We discovered that, in parallel with learning, populations of neurons emerge in cortical, thalamic, and extrathalamic regions of the somatosensory system that encode task-relevant stimulus features via the presence or absence of bursts. These burst-coder neurons (BCNs) increase in number with task proficiency and exhibit burstiness that scales with stimulus valence rather than physical stimulus identity. Notably, BCNs consistently track stimulus-outcome associations—even after multiple rule switches—by inverting their burst encoding of the physical stimuli, indicating that burst coding is driven by outcome associations rather than by inherent stimulus properties. Although burst coding emerges throughout the thalamocortical system, only cortical units retain significant burst coding after devaluation, while other regions lose their discriminative burst patterns. Furthermore, decoding of stimulus properties and behavior achieves maximal accuracy when bursts or BCNs are used as input. Overall, these results provide direct experimental evidence linking neuronal bursting to learning, supporting a novel perspective of bursts as context encoders and teaching signals.

## Main

The ability of neurons to fire high-frequency action potentials (bursts) is found across different brain regions and species^1–3^. Bursts have been intensively studied in the context of their cell-intrinsic mechanisms leading to seminal discoveries of burst-generating mechanisms in the thalamus^4^ and cellular mechanisms for input coupling across cortical layers^5^. To date, bursts are functionally understood as carriers of stimulus-related information, such as sensory features, perception or brain state^6–10^. On the other hand, high-frequency action potentials are well-documented to drive synaptic plasticity^11–17^, but the relationship between neuronal bursts and learning remains largely unexplored. Recent simulations and mathematical analyses support the idea that bursts may act as learning signals to instruct synaptic plasticity without affecting the processing of sensory information^18^. Despite the strong evidence linking synaptic plasticity to high-frequency firing, direct experimental evidence connecting bursts to learning is lacking.

Here, we employ a novel whisker discrimination learning task^19^ in freely behaving mice, combined with multi-site recordings of unit activity in four key regions of the whisker thalamocortical system, to directly investigate a potential link between neuronal bursts and learning.

### Aperture discrimination learning during different task rules in mice

In this go/no-go operant learning task we trained mice to associate apertures of different widths with a respective reward or punishment (Fig. 1a). Food-restricted mice were freely roaming on a linear platform with one lick-port at each end of the platform. To reach the lick-ports, mice had to pass through two-winged motorized apertures, which they touch with their whiskers. Licking was registered at the lick-ports and triggered either a reward (condensed milk) or a punishment (white noise, 120 dB, random duration between 1 to 3 seconds). In the initial rule, we trained mice to associate the wide aperture with reward and the narrow aperture with punishment. Mice learned to discriminate between the two apertures (d’ above 1.65) after an average of 13.5 sessions (± 4.4 SD, n = 6 mice) or 388 trials (± 53 SD, n = 6), corresponding to seven days of training (Fig. 1b, Extended Data Fig. 1 & Supplementary Video 1-3). We validated the whisker-dependency of the task by removing their whiskers (comparing the sessions before and after whisker removal with a performance of 2.66±0.41 SD and 0.60±0.47 SD respectively, p-value 2.0795e-08, two-sided paired t-test, n = 14) (Fig. 1c) and by numbing their whisker pads with a topical anaesthetic (lidocaine cream)^19^, both of which abolished task performance. We trained mice in four consecutive training stages with distinct stimulus-outcome contingencies (Fig. 1d). The introduction of a third aperture, which provided neither reward nor punishment (henceforth referred to as the “neutral aperture”), did not disrupt overall task performance. Introducing a rule reversal abolished performance and led to subsequent re-learning of the reversed rule, albeit over a significantly extended period (1077±330 trials, p-value compared to initial learning speed 0.03125, Wilcoxon signed rank test, n = 6, Extended Data Fig. 1). Degrading the aperture-outcome contingencies by randomizing reward and punishment presentations resulted in a complete loss of discrimination ability, yielding an average performance of d prime of -0.02 (±0.20 SD, n = 40 sessions). The ability to train mice in multiple rule settings in this paradigm allowed us in the following to examine how stimulus-outcome associations are encoded in the brain to drive learning.

**Fig. 1.**
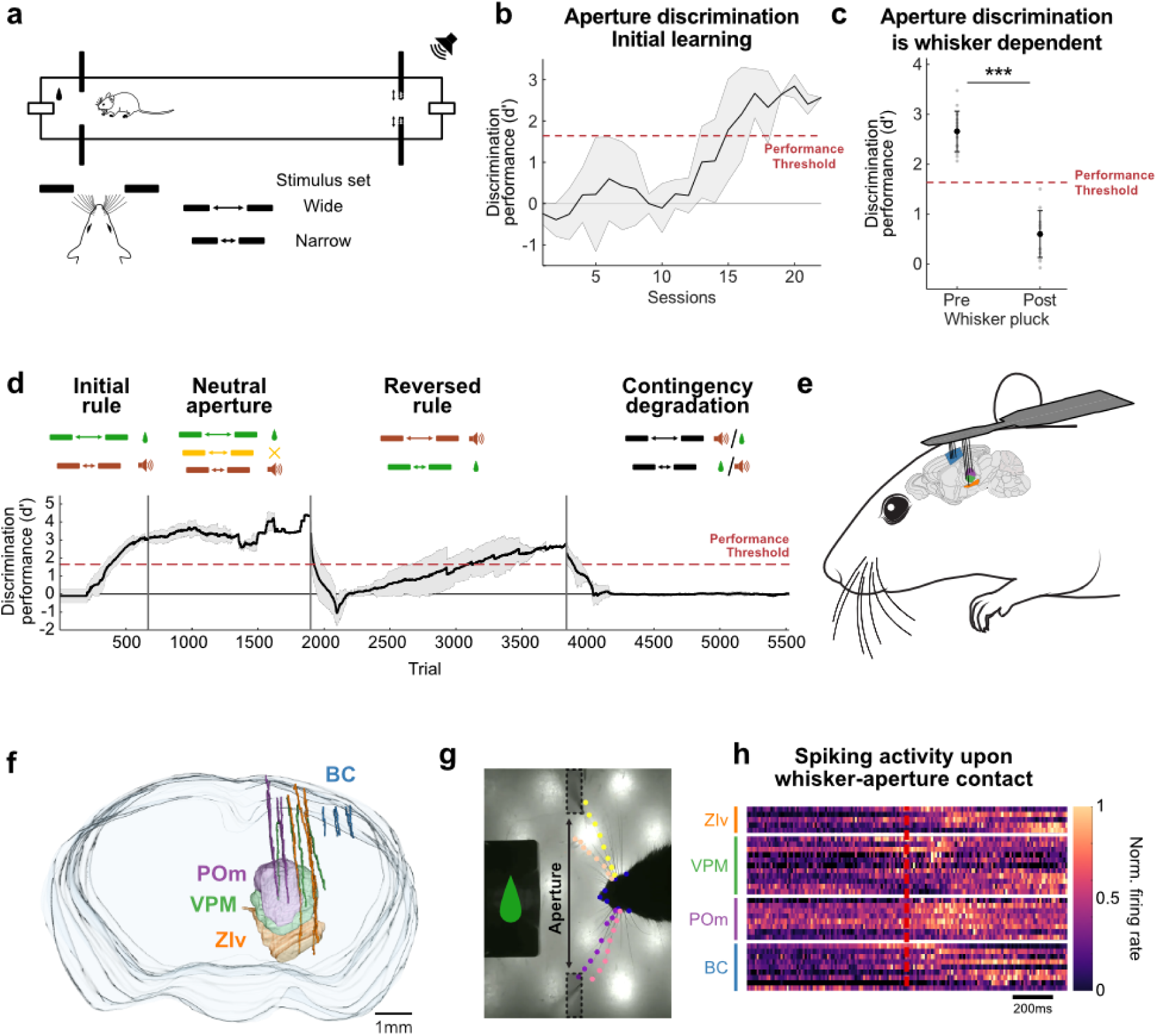
Whisker-dependent discrimination task to study neuronal dynamics during learning in freely moving mice. **a,** Schematic of the whisker-dependent aperture discrimination paradigm, where the wide aperture is rewarded (“Hit”) and the narrow aperture is punished (“Failure”). **b,** Learning curve for the initial rule from six mice (average ± std). Expert-level discrimination performance (d’ ≥ 1.65, red dashed line) was achieved after 13.5 (± 4.4) sessions. **c,** Discrimination performance before (Pre) and after (Post) whisker removal (p = 2.0795e-08, two-tailed paired t-test, n = 6). **d,** Performance across learning stages: initial learning, neutral aperture, reversed rule, and degradation. Each stage’s rules are indicated at the top, with average trial performance (black line) and standard deviation (shaded area). **e,** Schematic of multi-site tetrode recordings: 16 tetrodes were implanted in BC (5), VPM (4), POm (4), and ZIv (3). **f,** Example tetrode track reconstruction from post-mortem histology. **g,** High-speed camera frame showing tracked whiskers (colored dotted lines) via DeepLabCut (20) to mark whisker-aperture touch events. **h,** Example spiking activity from BC, VPM, POm, and ZI in response to whisker-aperture touches (red line), normalized from 0 to 1.

### Burst-coding neurons (BCNs) encode the aperture width through the presence or absence of bursts

We investigated four key brain regions implicated in somatosensory processing by recording multi-site unit activity (Fig. 1e) in the barrel cortex (BC), the ventral posteromedial nucleus (VPM), the posterior medial nucleus of the thalamus (POm), and in the zona incerta ventralis (ZIv). Post-mortem tetrode track reconstructions validated tetrode localization (Fig. 1f), and only data from correctly placed tetrodes were included in the analysis. The time points of the whisker-aperture interactions were extracted from the high-speed videos using DeepLabCut^20^ (Fig. 1g, h). In the following, we present the results from a cohort of six mice which went through the four training stages (Fig. 1d) while their unit activity was recorded in BC, VPM, POm, and ZIv. We split units according to waveform analysis into regular-(RS) and fast-spiking (FS) units (Extended Data Fig. 2). We restricted the following analyses on RS units in BC, VPM and POm and RS/FS units in ZIv, unless otherwise stated.

First, we quantified the proportion of units responsive to whisker touch and locomotion across all recording sites (see Methods). We observed a substantial increase in the proportion of whisker touch responsive units, from 12.5% (±3.2%) in the initial third to 52.2% (±2.5%) in the final third of the learning phase. In contrast, the proportion of locomotion-encoding units remained relatively constant throughout learning at around 10% (Fig. 2a). Looking at individual recording sites, we found robust whisker touch responses in each of the four recorded regions (Fig. 2b). Whisker-touch responses consisted of a mix of both tonic and burst spikes in all four regions. Baseline burst fractions varied among the regions and peaked shortly after touch onset (Extended Data Fig. 3a).

**Fig. 2.**
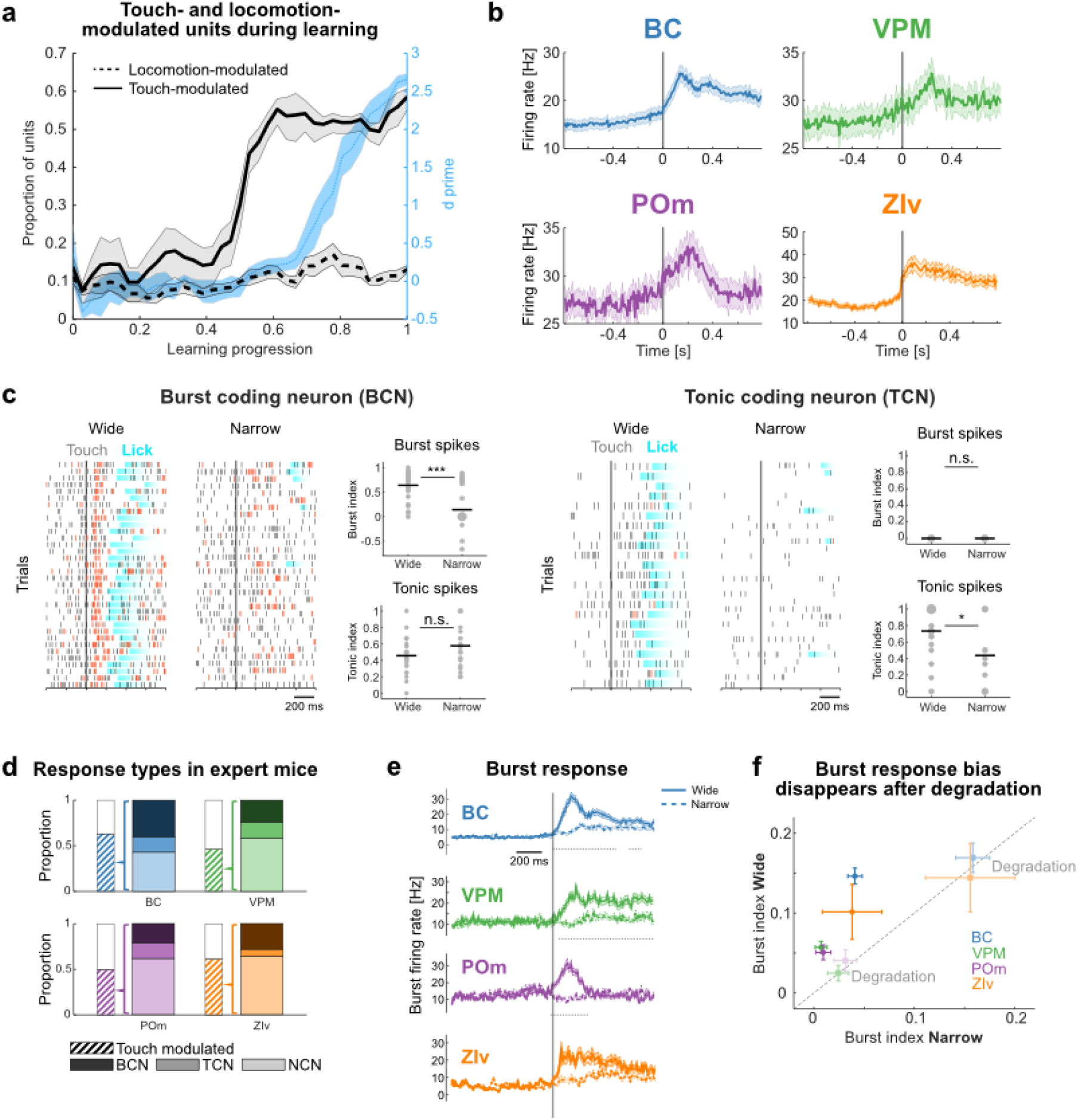
Burst encoding of aperture width during task performance. **a,** Proportion of whisker-touch-(solid line) and locomotion-modulated (dashed line) units across learning. Whisker-touch modulation is defined by significant responses to the apertures (two-sided Wilcoxon signed rank test, based on unit firing rates), and locomotion-modulated units were identified by stepwise multiple linear regression. Discrimination performance (d′) is indicated with a blue line. **b,** Mean population firing rate (shaded areas show SEM) in response to aperture whisker-touches in expert sessions (d′ ≥ 1.65). **c,** Spike rasters and burst and tonic indices for example BCN (left) and TCN (right). Burst spikes are marked in red, lick events in blue. **d,** Proportion of response types (BCNs, TCNs, and NCNs) in expert mice across regions (for exact values see Extended Data Table 1). **e,** Burst firing rates of BCNs in response to wide (solid) and narrow (dashed) apertures. Significant differences (α = 0.05) are indicated with dotted lines. **f,** Mean burst indices per region, split between wide and narrow apertures, with SEM as error bars. Dotted lines indicate equal burstiness for wide and narrow apertures. The burst bias in expert sessions (circles) disappears during the degradation stage (pale squares). n.s.: p > 0.05, *: p ≤ 0.05, **: p ≤ 0.01 and ***: p ≤ 0.001; **c:** two-tailed Wilcoxon signed rank test, **e:** two-tailed Welch’s t-test. For exact p values see Extended Data Table 1. **a,b,d-f:** data pooled from n= 6 mice.

We then asked if the aperture width is encoded in the spiking response profiles. By inspecting response profiles of individual units when mice encountered either the wide or narrow aperture, we identified a subset of units which exhibited significantly higher bursting activity for the wide aperture compared to narrow aperture (Fig. 2c, left). We refer to these units, which encode the aperture state through the presence or absence of bursts, as burst-coding neurons (BCNs). A different subset of units also encoded the aperture state, but with tonic spikes (Fig. 2c, right), henceforth referred to as tonic-coding neurons (TCNs). The remainder of touch-responsive units did not respond differently to the wide and narrow aperture and these are referred to as non-coding neurons (NCNs). Notably, we found that these three response types (BCNs, TCNs, NCNs) are present in all four recorded brain regions, with BCNs being the largest subset among the aperture coding units (Fig. 2d).

When we compared the burst-firing rate of BCNs between the wide and narrow apertures in a time-resolved manner the burst-coding of the aperture state became even more apparent. Across all recorded brain regions, wide-aperture-contacts consistently elicited higher burst-firing rates than narrow-aperture-contacts in BCNs but not in TCNs (Fig. 2e for BCNs, Extended Data Fig. 3b for TCNs). Notably, the POm exhibited only transient activation upon aperture-contact (Fig. 2e), while other brain areas showed prolonged activation after aperture-contact. Finally, we tested whether burst-coding is maintained after degrading the contingencies between stimulus and outcome during a degradation stage during which the aperture states and outcomes were randomized in each trial (50% chance for “wide” or “narrow” and reward or punishment, respectively) (Fig. 1d). We found that burst-coding disappeared following the degradation stage (Fig. 2f).

### BCNs track stimulus-outcome associations even after multiple rule reversals by inverting burst-encoding of the physical stimuli

An important consideration for the burst coding is whether BCNs encode the aperture’s physical properties (width) or their contextual parameters (outcome). We surmised that a mere encoding of the physical parameters would require the BCNs to retain their code (wide = burst, narrow = no burst), when the aperture context is changed during the reversed rule stage (Fig. 1d). We observed the opposite in that BCNs inverted their burst encoding of the physical stimuli when comparing narrow/wide encoding in expert sessions of the initial and reversed rule (Fig. 3a shows BC units, for VPM, POm, ZIv, see Extended Data Fig. 4). To explore how flexible the burst-code inversion is, we trained mice in an additional rule reversal stage following the reversed rule stage, where we restored the initial rule set (wide = rewarded, narrow = punished). After relearning, BCNs had again inverted their burst encoding back to the initial burst code (wide = burst, narrow = no burst). These results demonstrate that burst coding is not primarily determined by the physical properties of the stimuli, but rather by the association between the stimulus and the outcome and that BCN-bursts track the rewarded stimulus through multiple rule-reversals by inverting burst-encoding.

**Fig. 3.**
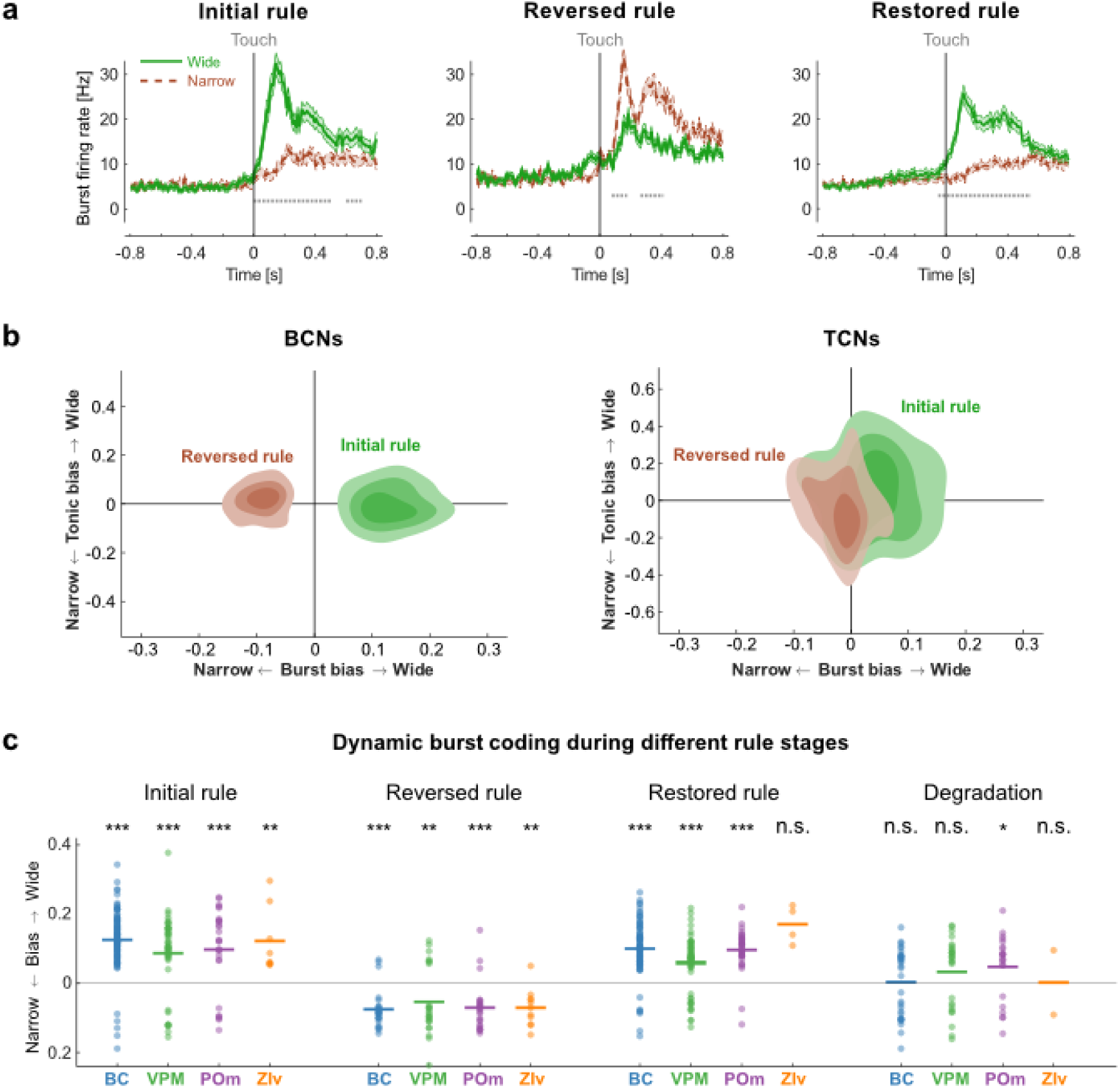
BCNs track stimulus-outcome associations even after multiple rule-reversals by inverting burst-encoding of the physical stimuli. **a,** Burst firing rates of BCNs in BC in response to wide (green) and narrow (maroon) aperture (represented as mean ± SEM, n = 6 mice). Statistically significant differences (α = 0.05) are indicated with dotted lines. For VPM, POm, ZIv, see Extended Data Fig. 4. **b,** Distribution of burst and tonic bias (see Methods for details) of BCNs (left) and TCNs (right), pooled from BC, VPM, POm, and ZIv (n = 6 mice). The initial rule is shown in green and the reversed rule in maroon. Contour lines correspond to the first (Q1), median (Q2), and third quartile (Q3) of units in the scatter plot. **c,** Burst bias of individual BCNs across the initial rule, reversed rule, restored rule, and degradation stage in BC, VPM, POm, and ZIv (n = 6 mice). n.s.: p > 0.05, *: p ≤ 0.05, **: p ≤ 0.01 and ***: p ≤ 0.001; **a:** two-tailed Welch’s t-test, **c:** two-tailed Wilcoxon signed rank test. For exact p values see Extended Data Table 1.

We explored the burst-code inversion in BCNs and TCNs in more detail by calculating burst- and tonic biases, which measure the occurrence of burst- and tonic spikes upon whisker touches of the wide or narrow aperture (see Methods). This revealed that (1.) BCNs employ burst coding and not tonic coding and that (2.) BCNs undergo a more pronounced code-inversion after rule reversal compared to TCNs which remained relatively stable (Fig. 3b shows units pooled from all regions, see Extended Data Fig. 5 for the individual brain regions).

A comparative longitudinal analysis of BCNs in BC, VPM, POm, and ZIv over the course of the different rule stages revealed that BCNs dynamically track the rewarded aperture with learning, independently of whether the rewarded aperture was wide or narrow. This analysis further shows that the burst bias is largely abolished by the degradation stage (Fig. 3c). Only the POm, retains significant burst coding, possibly due to the brevity of the degradation period.

### Burst coding scales with stimulus valence and predicts licking behavior

In the next training stage, we introduced a third aperture that was intermediate between the narrow and wide aperture and was neither rewarded nor punished (hence referred to as “neutral”). We observed that the lick rates for the intermediate aperture fell between those of the rewarded and punished apertures, indicating a possible scaling of lick behavior with stimulus valence (Fig. 4a). Similarly, the burst firing responses to the neutral aperture significantly differed from the others and was positioned between those of the rewarded and punished apertures in all four recording regions (Fig. 4b), supporting the notion that burstiness scales with stimulus valence.

**Fig. 4.**
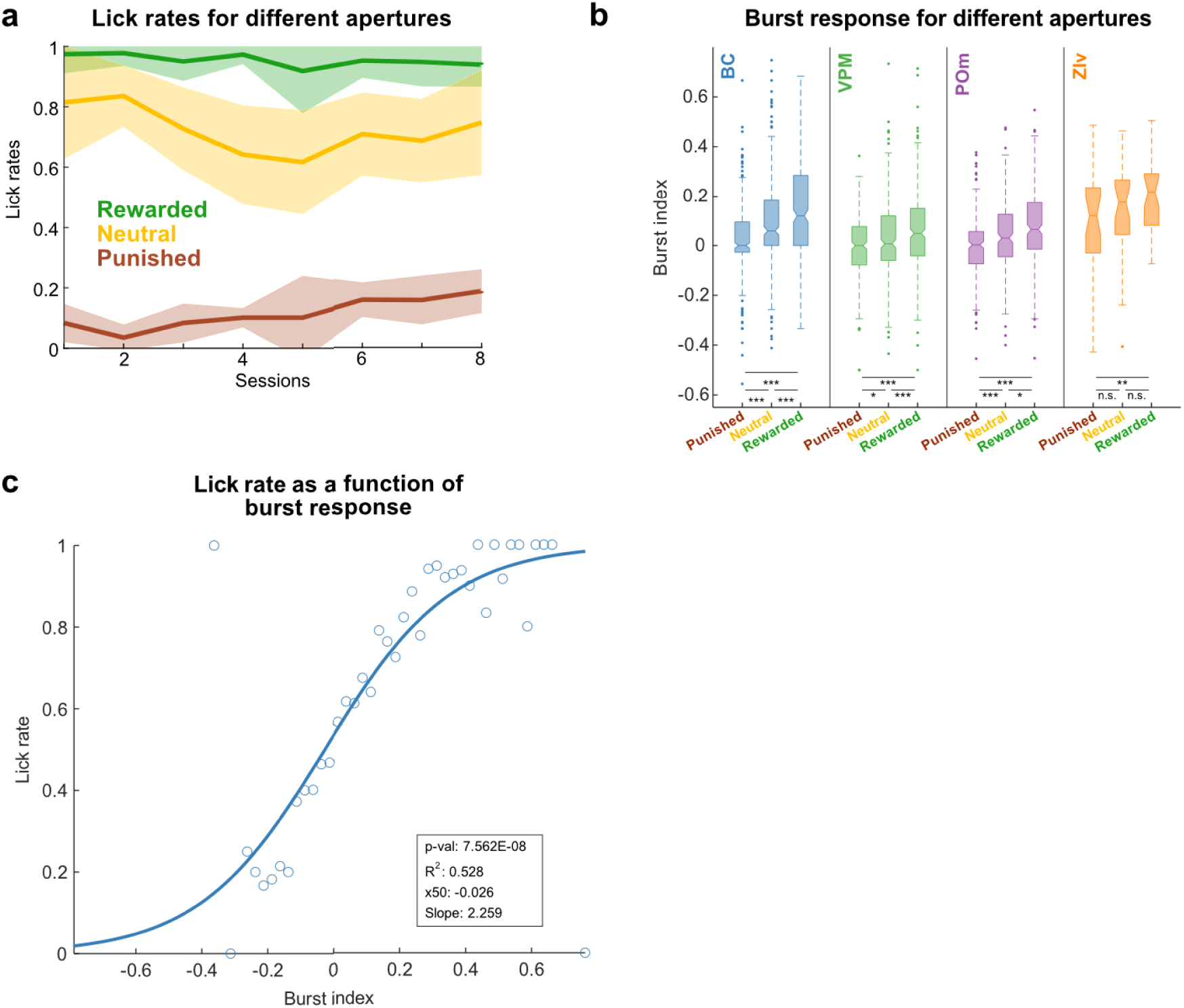
Burstiness and lick behavior scale with stimulus valence. **a,** Mean lick rates (SEM as shaded regions) across sessions of the neutral aperture stage. The three different apertures are represented separately (narrow/punished in red, intermediate/neutral in yellow, and wide/rewarded in green). **b,** Box plots comparing the burst indices in response to the different aperture types for each region (BC, VPM, POm, and ZIv) (see Methods). **c,** Lick rate as a function of burst bias in BC, considering, go, no-go as well as neutral trials. Properties of the logistic regression: R2: 0.528, p: 7.562e-08, inflection point (x50): -0.026, slope: 2.259. n.s.: p > 0.05, *: p ≤ 0.05, **: p ≤ 0.01 and ***: p ≤ 0.001; **b:** two-tailed Wilcoxon signed rank test. For exact p values see Extended Data Table 1. Data pooled from n = 6 mice.

Furthermore, we explored whether the mouse’s decision to lick in a given trial is predicted by the strength of the burst coding during aperture sampling. Notably, the likelihood of a lick event (calculated as the proportion of trials with a lick outcome for a given burst bias) increased continuously with the burst bias (Fig. 4c), as captured by the logistic fit, which significantly predicts lick rate with good accuracy (R² = 0.528, p = 7.562e-08). The inflection point of the fit, occurring near zero burst bias (x50 = -0.026), suggests that modulation of bursting (either suppression or enhancement) could serve as an effective neural signal to inhibit or drive reward-seeking behavior. This underscores the close relationship between burst coding and behavioral outcomes.

### Burst coding emerges with learning and is correlated with task performance

We next investigated the relationship between task proficiency and burst coding by grouping the sessions of the six mice into performance categories and computing the strength and direction of aperture burst coding for each category. As performance improved, a burst bias towards the rewarded aperture emerged, supporting a direct link between task proficiency, and burst coding (Fig. 5a shows units pooled from all regions, see Extended Data Fig. 6 for individual brain regions). Upon rule reversal, the burst bias was initially inverted, concomitantly with inverted performance (negative d’), reflecting a carryover from prior performance under the initial rules. With increased proficiency in the reversed rule stage, the burst bias shifted to favor the – now-rewarded – narrow aperture (Fig. 5a).

**Fig. 5.**
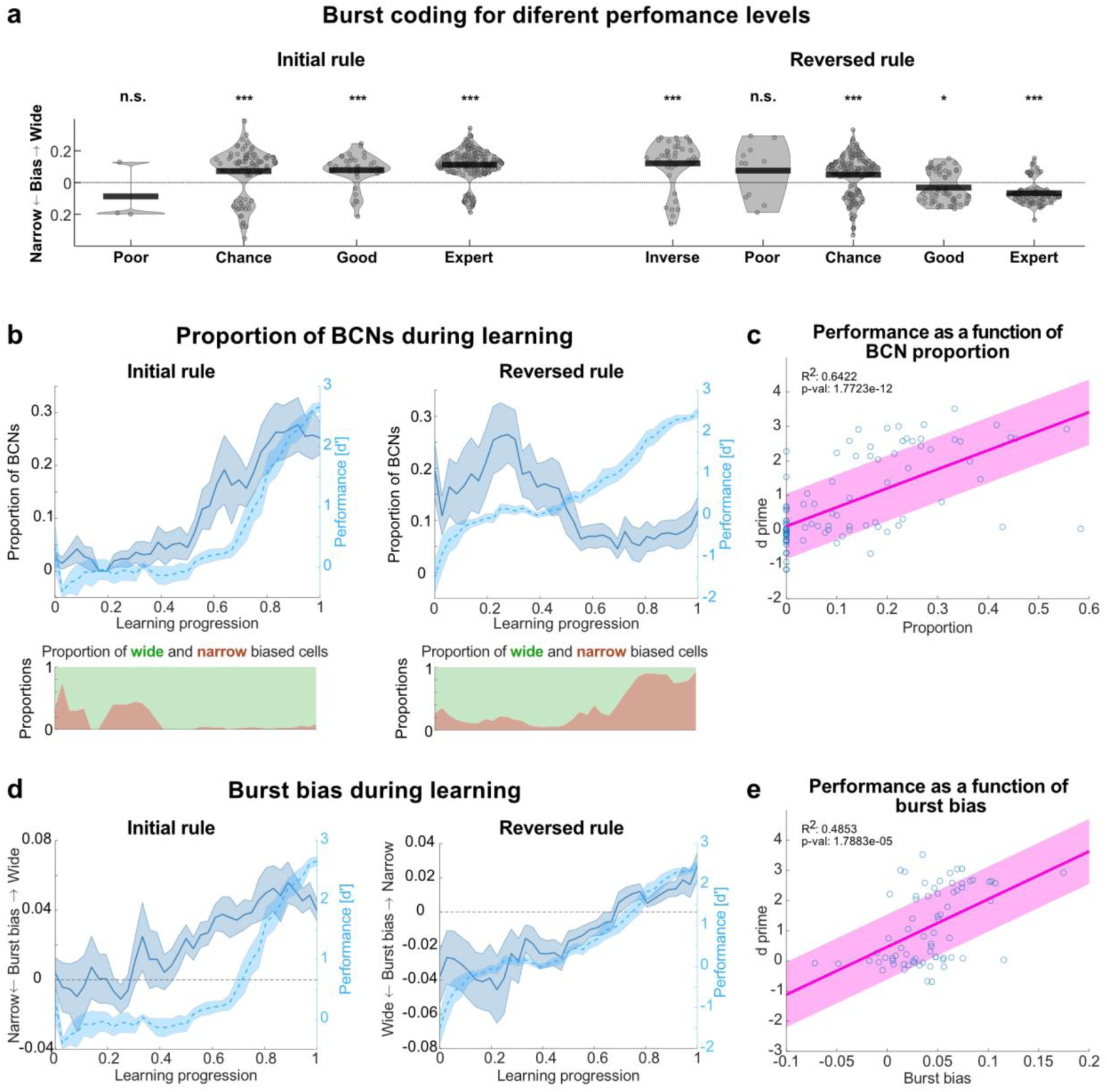
Burst encoding emerges with learning. **a,** Violin plots of burst bias in pooled BCNs across BC, VPM, POm, and ZIv regions, shown for different performance levels and learning stages. Performance levels are defined as inverse (d’ ≤ -1.65), poor (-1.65 < d’ < -0.5), chance (-0.5 ≤ d’ ≤ 0.5), good (0.5 < d’ < 1.65), and expert (d’ ≥ 1.65). **b,** Line graphs showing the average proportion of BCNs in BC (shaded SEM) as learning progresses, with mean performance (d′) indicated by a dotted line. Right panel shows data under the reversed rule, with proportions of wide- and narrow-biased BCNs below. **c,** Positive correlation between performance and BCN proportion (R^2^: 0.6422, p: 1.7723e-12). **d,** Average burst bias in BC across animals over learning, with bias between narrow and wide apertures shown for the initial rule (left) and reversed rule (right). Average d′ is indicated by the dotted line. **e,** Session performance correlates with the average burst bias per session (R^2^: 0.4853, p: 1.7883e-05). For VPM, POm, ZIv, see Extended Data Fig. 7 and 8. n.s.: p > 0.05, *: p ≤ 0.05, **: p ≤ 0.01 and ***: p ≤ 0.001; **a**: two-tailed Wilcoxon signed rank test. For exact p values see Extended Data Table 1. Data pooled from n = 6 mice.

We next asked whether burst coding emerges during learning, by calculating the proportion of BCNs and their burst bias as a function of learning progression. In the initial learning stage, the proportion of BCNs in the BC increased in parallel with improving performance (Fig. 5b, left). After rule reversal, this proportion gradually declined, primarily attributed to a reduction in BCNs coding for the wide aperture, in favor of those coding for the narrow aperture (Fig. 5b, right). Notably, the proportion of BCNs shows a strong positive correlation with performance (R²: 0.6422, p: 1.7723e-12), highlighting the critical role of BCNs during learning and performance optimization (Fig. 5c). Similarly, the average burst bias in BC across animals evolved alongside learning, progressively favoring the rewarded aperture (Fig. 5d, left). Notably, during the rule reversal, we observed an inversion in the average burst bias, reflecting a shift in neural burst coding patterns (Fig. 5d, right). The average performance per session was positively correlated with the burst bias exhibited within that session (R²: 0.4853, p: 1.7883e-05), indicating that the strength of burst coding is closely linked to the progression of learning (Fig. 5e). Similar results were observed in all four recorded regions (for VPM, POm, ZIv, see Extended Data Figs. 7, 8).

### BCNs predict aperture states better than other touch-modulated units

Next, we addressed whether and to which extend aperture widths can be decoded from BCN activity by training a classifier to predict which aperture was sampled based on unit spike trains from the different recording sites (see Methods). The decoding accuracy of the classifier increased progressively with the number of units used as input. Notably, the performance of the classifier was consistently higher when trained on BCNs compared to other touch-modulated units (Fig. 6a-d). To achieve a decoding accuracy of 80% or higher, the number of units required was significantly lower for the BCN group (7 units for BC, 25 for VPM, and 9 for POm) compared to the touch-modulated group including BCNs (18 units for both BC, 76 units for VPM, and 31 for POm) and excluding BCNs (46 units for BC, more than 80 for VPM, and 77 for POm) (two-tailed paired t-test with an α = 0.05). This demonstrates the superior predictive power of BCNs in decoding sensory information and trial outcomes.

**Fig. 6.**
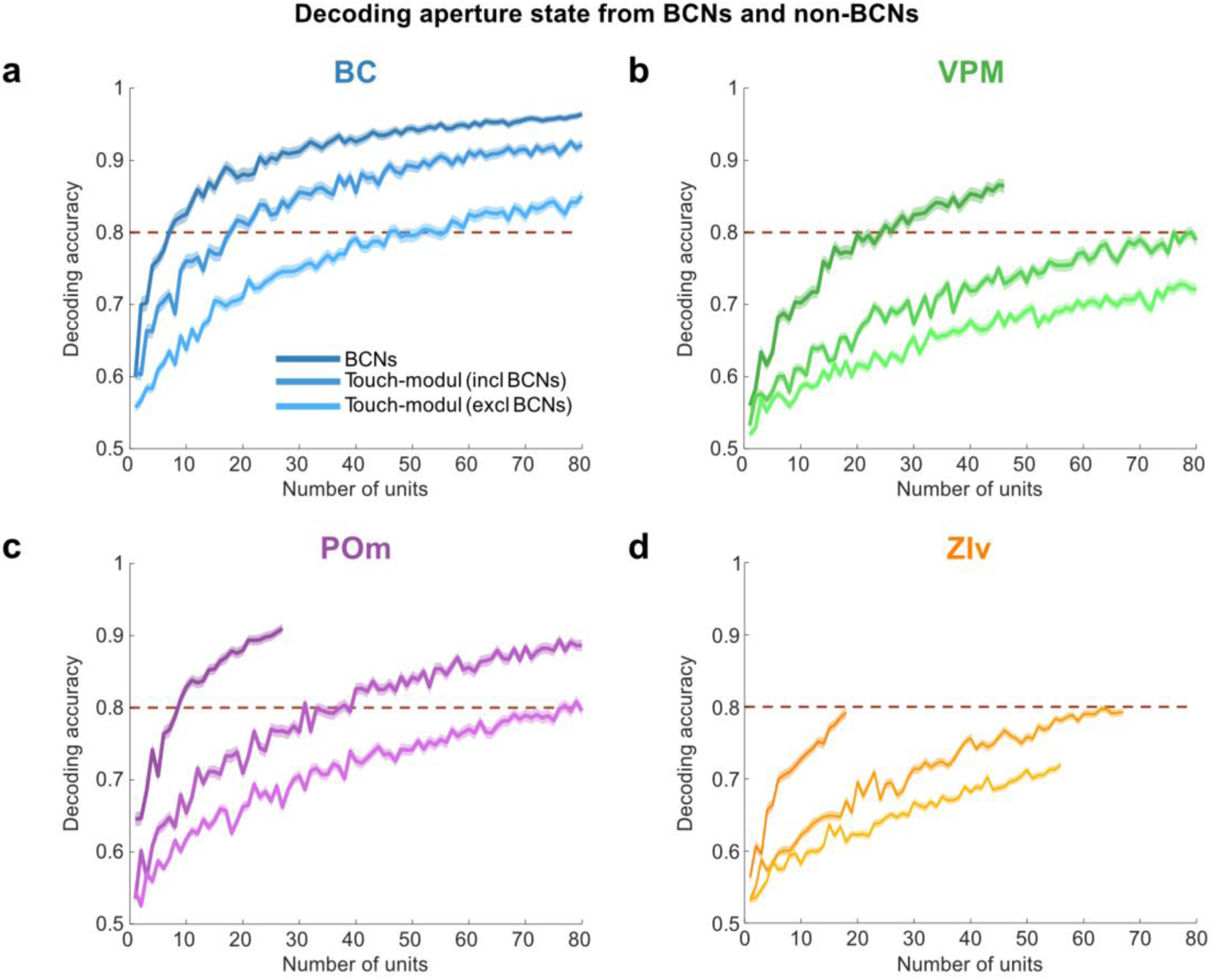
BCNs predict aperture states better than other touch-modulated units. **a-d,** Average decoding accuracy within a 400 ms window following whisker touch, plotted as a function of the number of units. The decoding accuracy curves are presented for BCN units alone, touch-modulated units including BCNs, and touch-modulated units excluding BCNs. The red dotted line marks a decoding accuracy threshold of 80%. Data is shown for the four recorded brain regions: BC (**a**), VPM (**b**), POm (**c**), and ZIv (**d**). Data pooled from n = 6 mice.

## Discussion

These results provide evidence for a relationship between learning and burst activity that emerges in burst-coding neurons (BCNs) across cortical, thalamic and extrathalamic regions. BCN units encode and track the stimulus context during different rule stages and burst coding largely diminishes during un-learning, as seen in the degradation stage.

The emergence of burst coding during learning may be an important mechanism to strengthen synaptic connections between neurons that form the memory engram for task execution. Ex-vivo work has demonstrated that burst firing modulates long-term synaptic coupling via calcium influx through NMDA receptor channels, a critical process for altering synaptic strength^14,21,22^. In this framework, sensory-evoked burst firing in BCNs could serve as the learning signal needed for the formation of stimulus-outcome association memories. Our observations of the steepest increase in BCN proportions during initial task learning, followed by the loss of burst coding during degradation, suggest that BCN bursts are particularly important during the learning phase. This relationship between burst firing and task performance aligns with previous research, which has shown that increases in burst activity correlate with improved detection probability and perceptual salience, both in the somatosensory cortex^9,23^ and the prefrontal cortex^24^.

The observed inversion of burst-coding following a rule reversal suggests that BCNs track the stimulus context rather than the stimulus’ physical features. As proposed by Payeur et al. (2021)^18^ for synaptic plasticity in hierarchical networks, the feedback responsible for carrying the credit must be assigned and maintained closely linear relative to the credit information. Our experimental results support this learning model, showing that burst responses are biased toward the rewarded stimulus and are modulated linearly according to valence, which is particularly evident when a neutral stimulus is introduced.

Our observation of distributed burst coding in all four recording sites in the somatosensory system, along with more localized findings in cortical layer 2/3 neurons of auditory cortex^25,26^, hints towards a universal principle of neural burst coding in learning and memory across different modalities.

## Methods

### Animals

All experimental procedures received approval from the local governing authority (Regierungspräsidium Karlsruhe, Germany, approval numbers: 35-9185.81/G-216/19) and were conducted in accordance with their ethical guidelines. The study involved adult male C57BL/6NRj mice (Janvier Labs, Le Genest-Saint-Isle, France) aged 8-10 weeks at the onset of training. The mice were individually housed in a ventilated Scantainer (Scantainer Classic, SCANBUR A/S, Karlslunde, Denmark) maintained on a 12-hour inverted light/dark cycle (lights off at 7:00 a.m. and on at 7:00 p.m.) with controlled temperature (22-25 °C) and humidity (40-60%). During behavioral training, which was conducted during day time, mice were subjected to a food restriction regimen, maintaining their body weight at 85-95% of their initial weight, with ad libitum access to water.

### Behavioral setup

The setup is described in detail in Heimburg et al. (2024)^19^ and a parts list, technical drawings and code can be found on Zenodo^27^. In brief, behavioral testing and recording were conducted on an elevated, dark-toned linear platform featuring lick ports at both ends. Mice traversed an adjustable aperture between two motorized wings to access the lick ports, with aperture widths varying trial-to-trial. Each lick port was equipped with a cannula connected to a syringe filled with condensed milk, and a speaker for white noise delivery. A piezo sensor on the cannula registered lick events that triggered either a reward (10 µL of condensed milk) or a punishment (120 dB noise, lasting 1-3 seconds). The setup was controlled via Syntalos^28^ (code available at https://github.com/bothlab/syntalos), which synchronized the timing of events (aperture state, reward/punishment, lick detection, and IR beam crossings), cameras, and electrophysiology.

### Behavioral paradigm

The paradigm is inspired by Krupa et al. (2001)^29^ and was recently adopted by us for mice as described in detail in Heimburg et al. (2024)^19^. In brief, mice were trained on a go/no-go whisker-dependent task on a linear track, learning to discriminate between apertures of different widths to obtain rewards and avoid punishments. Training sessions lasted 15 minutes, conducted twice daily with at least a two-hour break between sessions. After sampling the aperture with their whiskers, mice decided to either lick or turn and run to the opposite end of the track. Aperture states were reset randomly at the beginning of each trial.

### Initial rule stage

Go (45 mm “wide”) and no-go (25 mm “narrow”) apertures were presented with an equal probability of 50 % in a random sequence. Licking upon the go (wide) aperture triggered a reward, and licking at the no-go aperture triggered a punishment.

### Neutral aperture stage

Reward and punishment contingencies remained as in the initial rule stage, but an additional “neutral” aperture (35 mm) was added. Licking upon the neutral aperture resulted in no outcome and no scoring. The neutral aperture stage lasted for 8 sessions.

### Reversed rule stage

Reward and punishment contingencies were inverted (i.e., the previously rewarded aperture was punished, and vice versa). The neutral aperture was not presented.

### Degradation stage

Aperture states and outcomes were randomized in each trial (50% chance for “wide” or “narrow” and reward or punishment, respectively) in order to degrade the apertures as an outcome predictor. The degradation stage lasted for 8 sessions.

### Data acquisition

Data acquisition during behavior was done with Syntalos^28^. An intan-module recorded all analog signals from the lick ports. The overview camera was connected to a recorder module that operated continuously throughout the session. The high-speed cameras and an event list were controlled by Syntalos via a custom-made Python script. The cameras were programmed to start recording if the animal approached the corresponding lick port, determined by the crossing of the corresponding light beam. The event list kept track of the triggered light beams and reported the state of the apertures. The list also noted if each trial was a success or failure.

### EIB manufacturing

Electronic Interface Boards (EIBs) were based on the design developed in Oettl et al. (2020)^30^. Details, such as technical drawings can be found in Heimburg et al. (2024)^27^. In brief, approximately 30 cm of tungsten wire (12.5 μm, Item # 100211, Tungsten 99.95% CS Wire, California Fine Wire Company) were twisted into a tetrode. Sixteen tetrodes, providing a total of 64 recording channels, were mounted into a custom designed EIB (produced by Multi-Circuit-Boards). The layout allowed for proper alignment of the tetrodes in the x-y plane. A custom-built scaffold was used to position the tips of the tetrodes before fixation on the EIB to ensure proper depth positioning of each tetrode during implantation (see Table 1). Tetrodes were then fixed to the EIB using UV-curable adhesive (3M™ Filtek™ Supreme XTE, 3M Deutschland GmbH, Neuss, Germany) and the opposite ends of the tetrode tungsten wires were then soldered onto the copper contacts of the EIB. Prior to implantation, the electrodes were suspended in a gold chloride solution (200 mg/dL gold chloride in deionized water, HT1004-100ML, Sigma-Aldrich, Inc., MO, USA) and gold plated to reduce their individual impedances to less than 100 kOhm. To test whether the channels conduct the signal well, a function generator (A365 Stimulus Isolator, World Precision Instruments, Inc., FL, USA) was connected to a basin with sodium chloride, in which the tetrodes were submerged. Only EIBs with ≥62 conducting channels were used for implantation.

**Table 1.** Tetrode targeting coordinates. A/P: antero-posterior, M/L: medio-lateral, D/V: dorso-ventral

| Tetrode | A/P [mm] | M/L [mm] | D/V with respect to dura [mm] |
| --- | --- | --- | --- |
| BC 1 | - 1.0 | 3.25 | - 0.4 |
| BC 2 | - 1.3 | 3.3 | - 0.3 |
| BC 3 | - 1.6 | 2.7 | - 0.85 |
| BC 4 | - 1.3 | 2.6 | - 0.85 |
| BC 5 | - 1.0 | 2.7 | - 0.85 |
| VPM 1 | - 1.95 | 1.95 | - 2.95 |
| VPM 2 | - 1.95 | 1.6 | - 2.95 |
| VPM 3 | - 1.75 | 1.75 | - 3.0 |
| VPM 4 | - 1.45 | 1.6 | - 2.95 |
| POm 1 | - 2.15 | 1.3 | - 2.75 |
| POm 2 | - 2.3 | 1.5 | - 2.75 |
| POm 3 | - 2.3 | 1.15 | - 2.8 |
| POm 4 | - 2.0 | 1.05 | - 2.85 |
| Zlv 1 | - 2.35 | 2.0 | - 3.7 |
| Zlv 2 | - 2.6 | 1.75 | - 3.7 |
| Zlv 3 | - 2.15 | 1.75 | - 3.75 |

### Surgical procedures for in vivo experiments

30 minutes prior to inducing general anesthesia, buprenorphine hydrochloride (Bupresol vet. Multidose 0,3 mg/ml, 10 ml, CP-Pharma Handelsgesellschaft mbH, Burgdorf, Germany) was administered subcutaneously at a dosage of 0.1 mg/kg of body weight. General anesthesia was induced using isoflurane (1-2% in oxygen, Isofluran Baxter, Baxter Deutschland GmbH, Germany), with careful monitoring and control of the eyelid reflex and pedal withdrawal reflex. Body temperature was maintained between 36-38 °C using heating pads. A corneal application of eye ointment (Bepanthen®, Bayer, Germany) and subcutaneous administration of saline solution (30 ml/kg of body weight, Fresenius Kabi Deutschland GmbH, Bad Homburg, Germany) were performed. The scalp was locally infiltrated with Lidocaine (Xylocaine® 1%, Aspen Pharma Trading Limited, Ireland) to anesthetize the surgical area. The animal was positioned in a non-traumatic stereotactic alignment apparatus (David Kopf Instruments, Tujunga, CA, USA). Small holes were drilled in the skull over the target recording sites using a dental drill (for exact coordinates see the Table 1 below; 78001 Microdrill, RWD Life Science, TX, USA). The EIB was then gradually lowered into the brain until the desired depth was reached. The EIB was attached to the skull using a cyanoacrylate-based adhesive (Super-bond, Sun Medical, Japan) and dental cement (Paladur®, Kulzer, Germany). Following the surgeries, the animals were placed on a heated surface within a pre-warmed enclosure before being returned to the ventilated cabinets (Scantainer Classic, SCANBUR A/S, Karlslunde, Denmark). Behavioral experiments started following an adequate period of recovery subsequent to EIB implantation.

The tetrode array containing 16 tetrodes (total of 64 recording channels), was implanted in the barrel cortex (BC; 5 tetrodes), the ventral posteromedial nucleus (VPM; 4 tetrodes), the posterior medial complex (POm; 4 tetrodes), and the ventral zona incerta (ZIv; 3 tetrodes). Coordinates are given with respect to bregma, according to^31^:

### Histology and identification of tetrode positions

After administering Ketamine-Xylazine through intraperitoneal injection (ketamine dosage: 120 mg/kg bw, CP-Pharma Handelsgesellschaft mbH, Burgdorf, Germany; xylazine dosage: 20 mg/kg bw, Xylavet^®^ 20 mg/ml, CP-Pharma Handelsgesellschaft mbH, Burgdorf, Germany), a transcardial perfusion with paraformaldehyde (PFA, 4% in phosphate-buffered saline [PBS]) was performed. The mouse’s head, along with the EIB implant, was post-fixed in PFA (4% in PBS) for a period of five days at 4°C. Subsequently, the EIB was removed, the brain was cut into 50 µm thick sections with a vibrating microtome (Thermo Scientific Microm HM 650V), and the tetrode traces were identified using a bright field microscope. Through the digital delineation of tetrode marks in each section, we were able to reconstruct the complete trajectory of each tetrode in three dimensions using the Amira software (v6.5, Thermo Fisher Scientific, Waltham, MA). Subsequently, all tetrodes located outside of the designated target regions were excluded from further analysis. Reconstruction files can be found on GitHub (https://github.com/GrohLab/Distributed-burst-activity-in-the-thalamocortical-system/tree/main/Data/Reconstructions).

### Data analysis

The behavioral data was analyzed using custom code developed in MATLAB R2023a (The MathWorks, Inc.), which is available for reference on GitHub (https://github.com/GrohLab/Distributed-burst-activity-in-the-thalamocortical-system). P-values below 0.05 were considered significant. Mean values were reported with standard deviations and median values with interquartile range (IQR).

### Statistical analysis of the learning process

Task performance in each session was measured using the discriminability index d-prime (d’), also known as Fisher discrimination index, as described previously^32,33^ defined as the difference between the z-scores of the hit rate and the false alarm rate: ***d′ = z(hit rate) — z(false alarm rate)***. Due to the constraints of the z-score transformation (which cannot handle proportions of 0 or 1), hit rates and false alarm rates of 0 are typically adjusted to 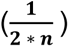 and rates of 1 are adjusted to (**1 −** 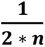), with ***n*** being the number of go (for hit rate adjustment), or no-go trials (for false alarm rate adjustment), respectively. A higher d’ indicates greater discriminability, and thus a greater discrimination performance of the mice. A d’ of 0 describes a discrimination performance at chance level. The criterion for achieving expert performance was established at a d-prime of 1.65, corresponding to a one-tailed significance level of α = 0.05.

### Extraction of behavioral data from videos

Spatial information of the mice was extracted from video recordings using DeepLabCut^20^. Following preprocessing with median background image subtraction, the DeepLabCut model was trained on a subset of manually annotated video frames. For the high-speed videos, these markers consisted of eight data points for two whiskers on each side from whisker base to tip, and markers for the mouse’s head contours. In overview videos, spatial positioning and velocity were estimated, based on anatomical landmarks on the mouse’s body and head.

### Electrophysiological data

The raw electrophysiological data was spike sorted with KiloSort 2.0. Subsequently, the sorted clusters were curated automatically using the Ecephys spike sorting algorithm provided by the Allen Brain Observatory (code available at https://github.com/alleninstitute/ecephys_spike_sorting). The algorithm includes noise templates to identify and exclude clusters exhibiting characteristics indicative of noise, such as irregular waveform shapes and inter-spike interval (ISI) histograms. The following quality metrics were then applied to further exclude any clusters indicative of multi-unit activity: isolation distance (computed from Mahalanobis distance) greater than 15 and ISI violation below 3%, with an ISI threshold of 1.5 milliseconds. The criteria were selected on the basis of previous studies^34–36^, where they have been demonstrated to effectively distinguish well-isolated single units from noise and multi-unit activity. The individual units were further classified into fast spiking (FS) and regular spiking (RS) cells, putatively representing interneurons and excitatory cells, respectively^37^. The threshold for differentiating RS from FS units was set to a trough-to-peak duration of 350 µs for cortex and zona incerta, and 300 µs for thalamic nuclei (Extended Data Fig. 2). These thresholds align with previous studies^38,39^ and consider the fact that extracellular waveforms in the thalamus are shorter than those in the cortex^38,39^. For further analysis, only RS units were selected for BC, VPM, and POm, while FS/RS units were selected for ZIv because of limited data in the literature in which FS/RS units in the ZI were associated with specific cell types (i.e., inhibitory / excitatory).

### Neural response metrics

#### Whisker-touch and locomotion responsiveness

Whisker touch responsiveness was defined by a significant increase in firing rates upon aperture touch within a 200 ms response window following aperture touch (to either of the two aperture states), analyzed using a two-sided Wilcoxon signed rank test. Locomotion-encoding units were identified through stepwise multiple linear regression analysis. This method is used to identify the significant predictors of a dependent variable by iteratively removing the least significant variables until all remaining variables are significant^40^.

### Bursts

Bursts are characterized as sequences of two or more spikes with anISI of ≤ 10 ms and a tail ISI of > 15 ms, in accordance with the criteria established previously^41,42^.

### Burst index

The burst index (*BI*) is calculated as the difference between the proportion of burst spikes (*P_bsp_*) in the response window (0 to 200 ms after touch) relative to the baseline window (600 to 400 ms before touch). *P_bsp_* is defined as the number of burst spikes (*N_bsp_*) over all spikes (*N_sp_*) in a given window. Values can range between -1 (maximal burst proportion in baseline with minimal burst proportion in response window) and 1. The formula is given by: 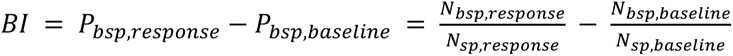

### Burst bias

The burst bias (*BB*) is derived from the burst indices of two different aperture conditions (*BI*_1_ and *BI*_2_). It is calculated as the difference between the two burst indices, divided by two. Values can range between -1 (maximal burst bias for aperture 2) and 1 (maximal burst bias for aperture 1). The formula is given by: 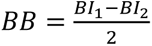

A significant burst bias classifies the unit as a burst-coding neuron (BCN), as determined by a Wilcoxon rank sum test.

### Burst fraction

The burst fraction (*f_b_*) is defined as the number of burst events (*N_be_*), disregarding the number of spikes within bursts, divided by the total event rate (*N_e_*), which includes both tonic spikes (*N_tsp_*) and burst events, as characterized in Friedenberger et al. (2023)^1^. The formula is given by: 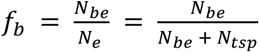

### Tonic index

The tonic index (*TI*) is calculated as the quotient of the number of tonic spikes (*N_tsp_*) in the response window (0 to 200 ms after touch) divided by the overall number of tonic spikes in the response and baseline windows (600 to 400 ms before touch) combined. Values can range between 0 (only tonic spiking in the baseline window) and 1 (only tonic spiking in the response window). The formula is given by: 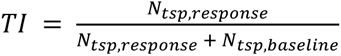

### Tonic bias

The tonic bias (*TB*) is derived from the tonic indices of two different aperture conditions (*TI*_1_ and *TI*_2_). It is calculated as the difference between the two tonic indices. Values can range between -1 (maximal tonic bias for condition 2) and 1 (maximal tonic bias for condition 1). The formula is given by: *TI* = *TI*_1_ − *TI*_2_

A significant tonic bias classifies the unit as a burst-coding neuron (TCN), as determined by a Wilcoxon rank sum test.

### Neural decoding analysis

The decoding of aperture states from neural spike time data was performed using the Neural Decoding Toolbox^43^. Classifier training was performed with an increasing number of units of either solely BCNs, touch-modulated units including BCNs, or touch-modulated units excluding BCNs. For each configuration, the dataset was divided into training (90% of labels) and test set (10% of labels). When insufficient units were available for partitioning, the test set was selected from the remaining units. Each set contained the spike traces of individual neurons for a given trial and the corresponding aperture labels. A support vector machine (SVM) classifier was trained using the LIBSVM software package. The classifier underwent 10-fold cross-validation, involving separate training and testing across different data partitions. To assess variability in decoding accuracy, each unit selection was bootstrapped 20 times. The mean decoding accuracy was calculated across a time window from trigger onset to 400 ms post-trigger onset.

## Data availability

The datasets generated and analyzed in this study will be made available in a persistent repository upon final publication.

## Code availability

The code used in this study is available in a GitHub repository (https://github.com/GrohLab/Distributed-burst-activity-in-the-thalamocortical-system/tree/main/Scripts).

## Acknowledgements

We thank Katharina Ziegler and Rebecca Mease for helpful inputs on the manuscript. We thank Wolfgang Kelsch and Max Scheller for teaching us the electronic interface board approach, Cornelius Schwarz for providing us with the lick ports, Lee Embray for building the behavioral apparatus and Matthias Klumpp for providing and helping with Syntalos. This work was supported by the German Research Foundation (DFG Grants GR3757/4-1 to AG). We acknowledge the data storage service SDS@hd and high-performance computing initiative bwHPC, supported by the Ministry of Science, Research and the Arts Baden-Württemberg (SDS@hd and bwHPC) and the German Research Foundation (DFG) through grants INST 35/1597-1 FUGG (bwHPC) and INST 35/1503-1 FUGG (SDS@hd). The funders had no role in study design, data collection and analysis, decision to publish, or preparation of the manuscript.

## Author information

Contributions Conceptualization: FH, AG Methodology: FH, AG Investigation: FH, JT, NMS Visualization: FH, JT Funding acquisition: AG Project administration: AG Supervision: FH, AG

Writing – original draft: FH, AG Writing – review & editing: FH, AG

## Extended data figures and tables

**Extended Data Table 1.**
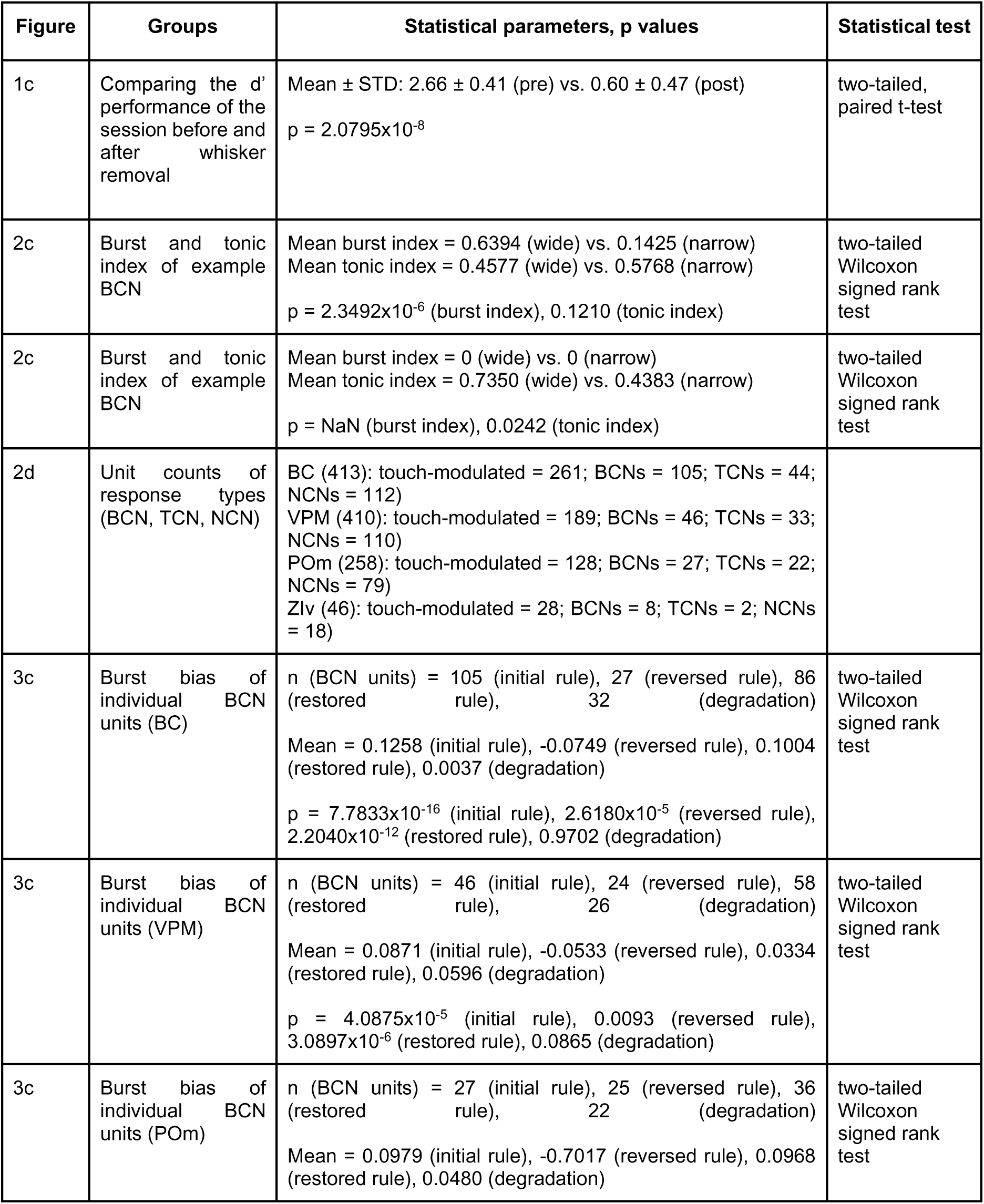
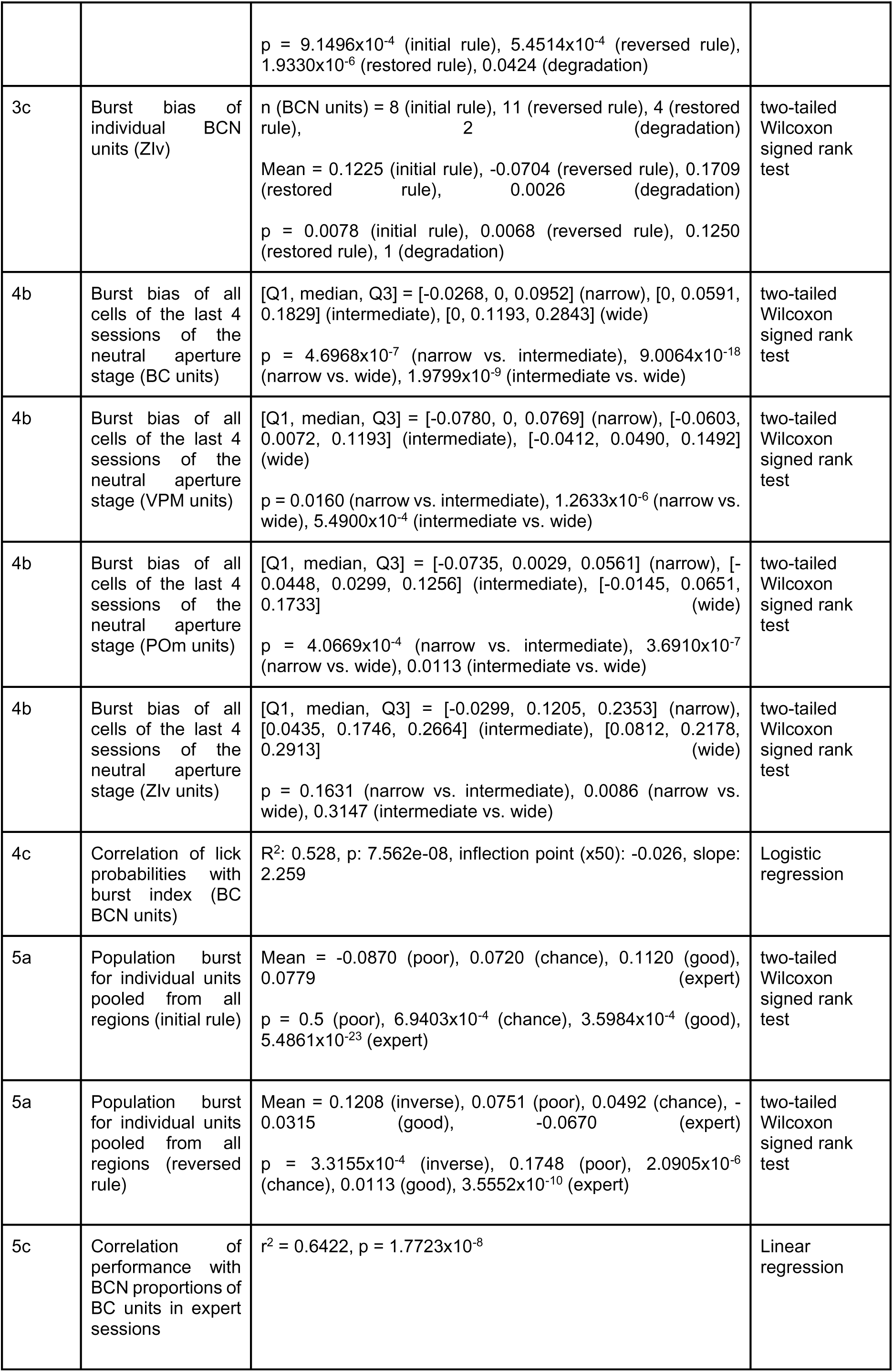
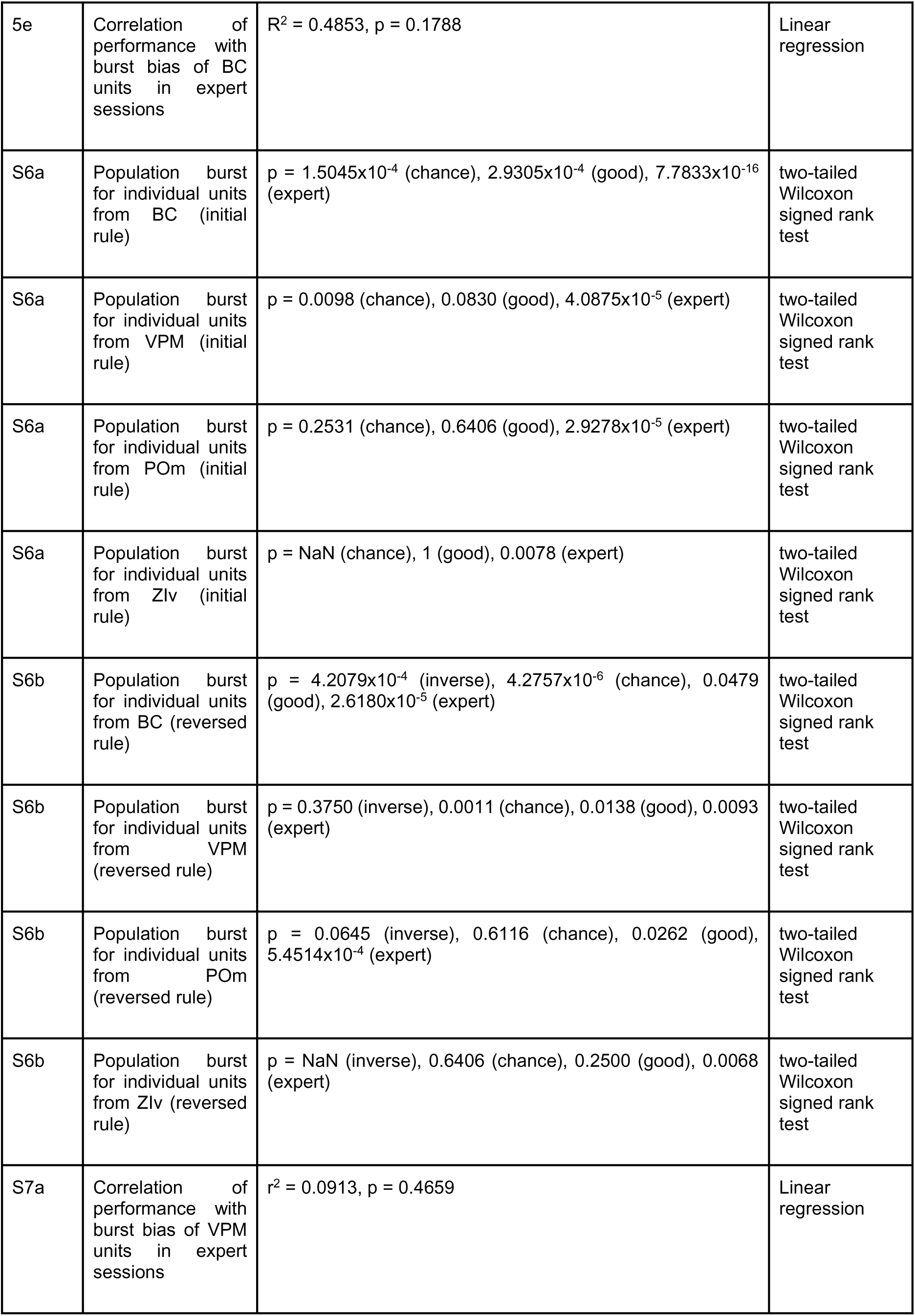
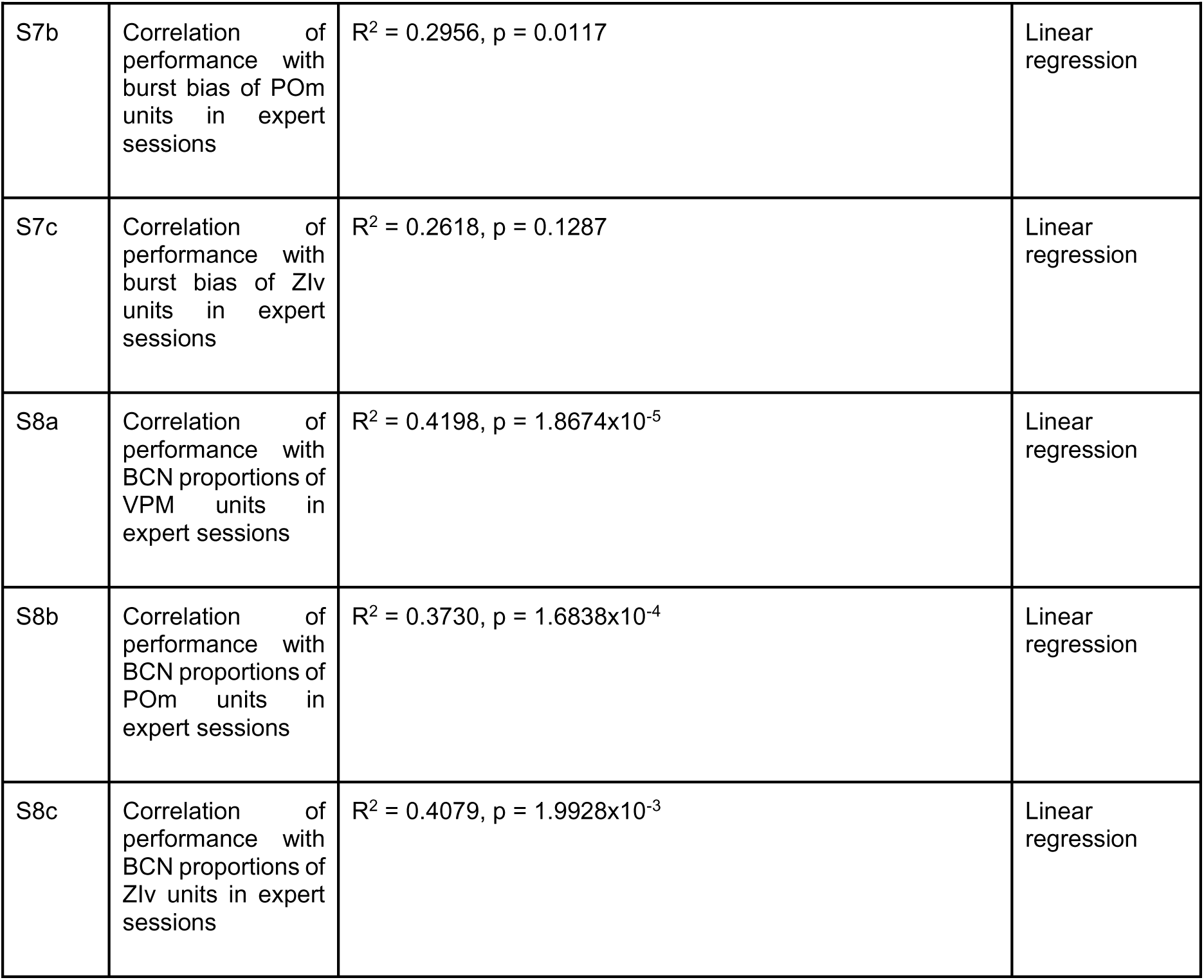
Description of statistical parameters.

**Extended Data Fig. 1.**
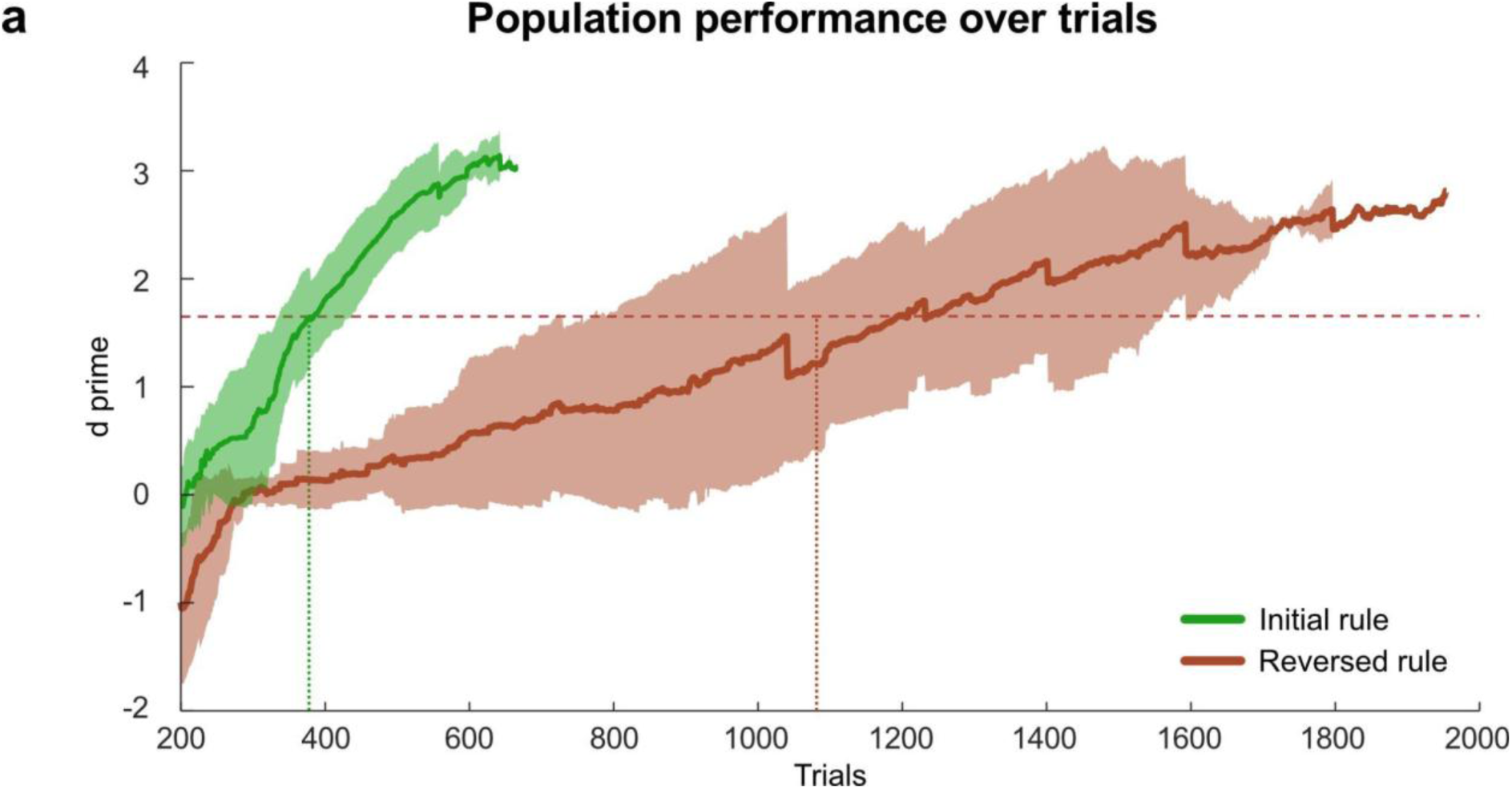
Population performance over trials. Mean d’ values are depicted as solid lines for both the initial rule (green) and reversed rule (orange), calculated using a running window of the preceding 200 trials. Shaded areas represent the standard deviation of the d’ values. The horizontal dashed line indicates the expert-level performance threshold (d’ = 1.65). Data was pooled from n = 6 mice. The dotted vertical lines mark the trials at which mice on average reached expert-level performance: 388±53 trials during the initial rule stage and 1077±330 trials during the reversed rule stage. Data pooled from n = 6 mice.

**Extended Data Fig. 2.**
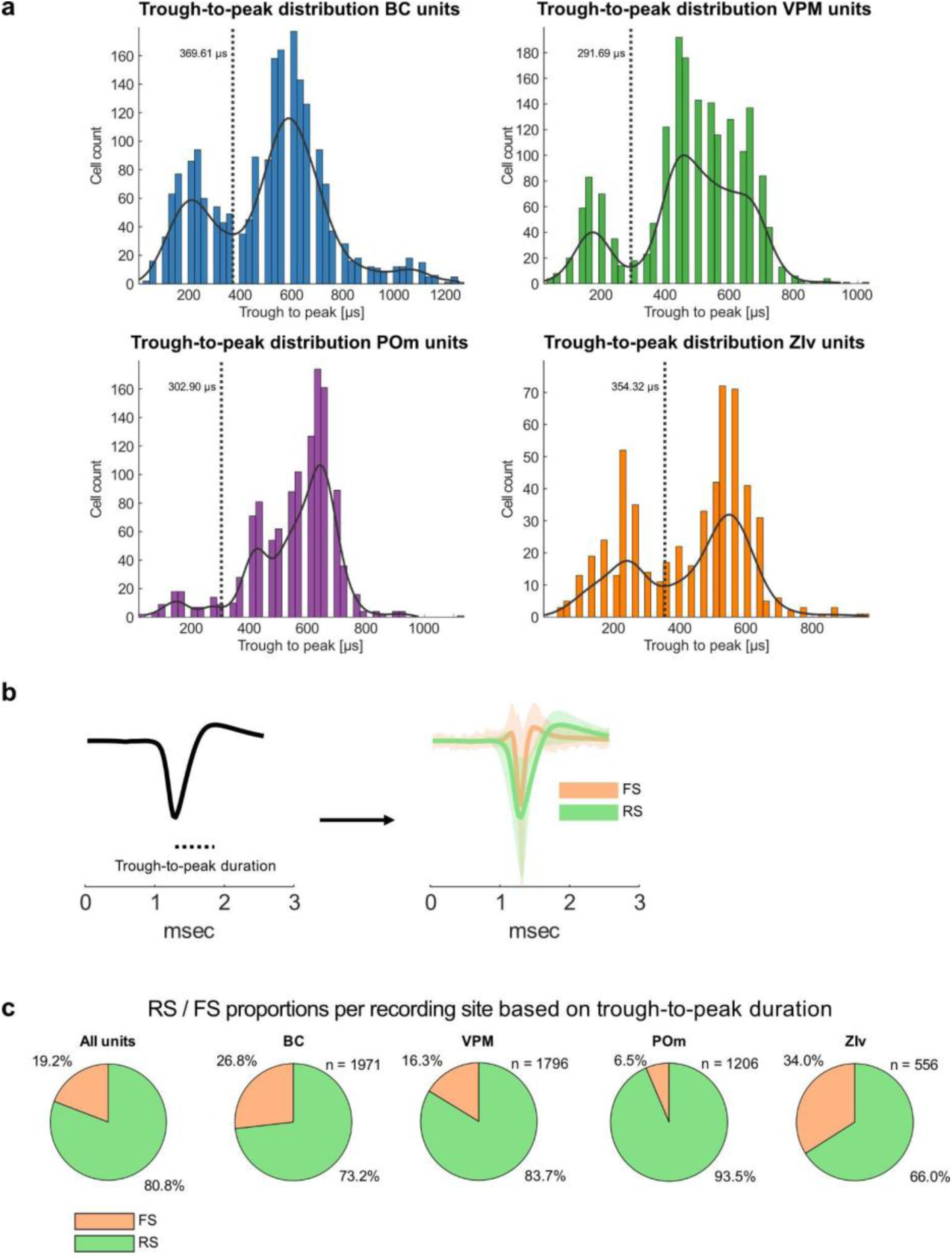
Unit waveform analysis across brain regions. **a,** Distribution of trough-to-peak durations per recorded brain region: BC (blue), VPM (green), POm (purple), and ZIv (orange), pooled from n = 6 mice. The dashed vertical lines represent the local minima that best distinguishes between regular-spiking (RS) and fast-spiking (FS) units in each region. **b,** Graphical illustration of the trough-to-peak duration metric, which is defined as the time interval between the trough and the second peak of the extracellular spike waveform. The overlaid waveforms (mean ± SEM) depict the two distinct subpopulations (RS and FS units). **c,** Breakdown of the proportion of RS (green) and FS (orange) units (putative excitatory- and inhibitory neurons, respectively ^37^) for each brain region pooled from n = 6 mice.

**Extended Data Fig. 3.**
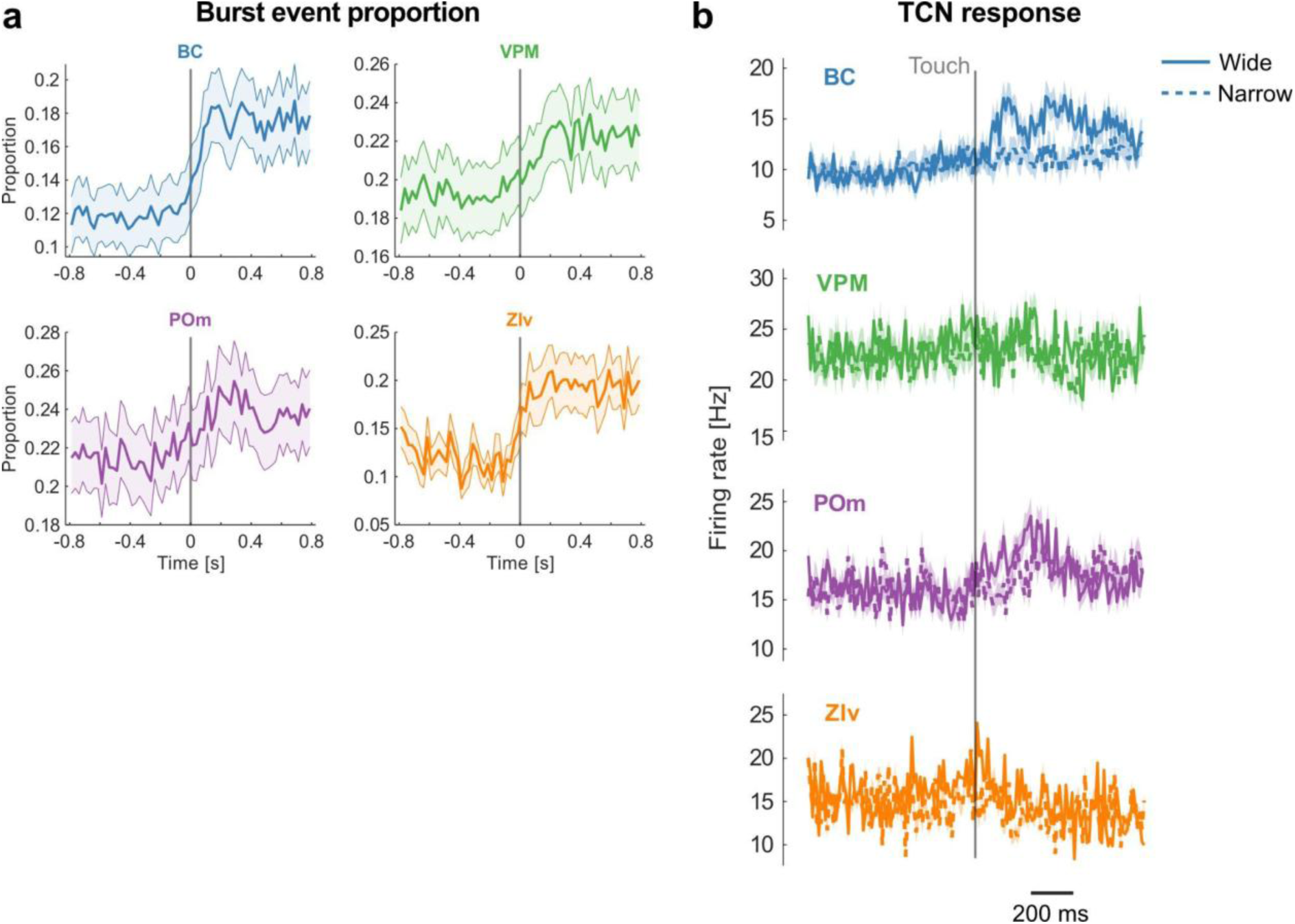
TCN whisker-touch responses and burst event proportions in expert animals of the initial rule. **a,** Burst fractions^1^ of touch-modulated units in response to whisker touches of the aperture (wide and narrow pooled). The mean burst event proportions develop from baseline values of 11.7 % (BC), 21.2 % (POm), 19.1 % (VPM), and 11.7 % (ZIv), to values after whisker touch of 17.2 % (BC, max. 18.7 %), 24.0 % (POm, max. 25.5 %), 21.8 % (VPM, max. 23.1 %), and 19.2 % (ZIv, max. 21.0 %). Data pooled from n = 6 mice. **b,** Tonic spike rates of TCNs in response to whisker touches of the wide (solid) or narrow (dashed) aperture. Firing rates in response to the two apertures do not differ significantly (two-tailed Welch’s t-test). Data pooled from n = 6 mice.

**Extended Data Fig. 4.**
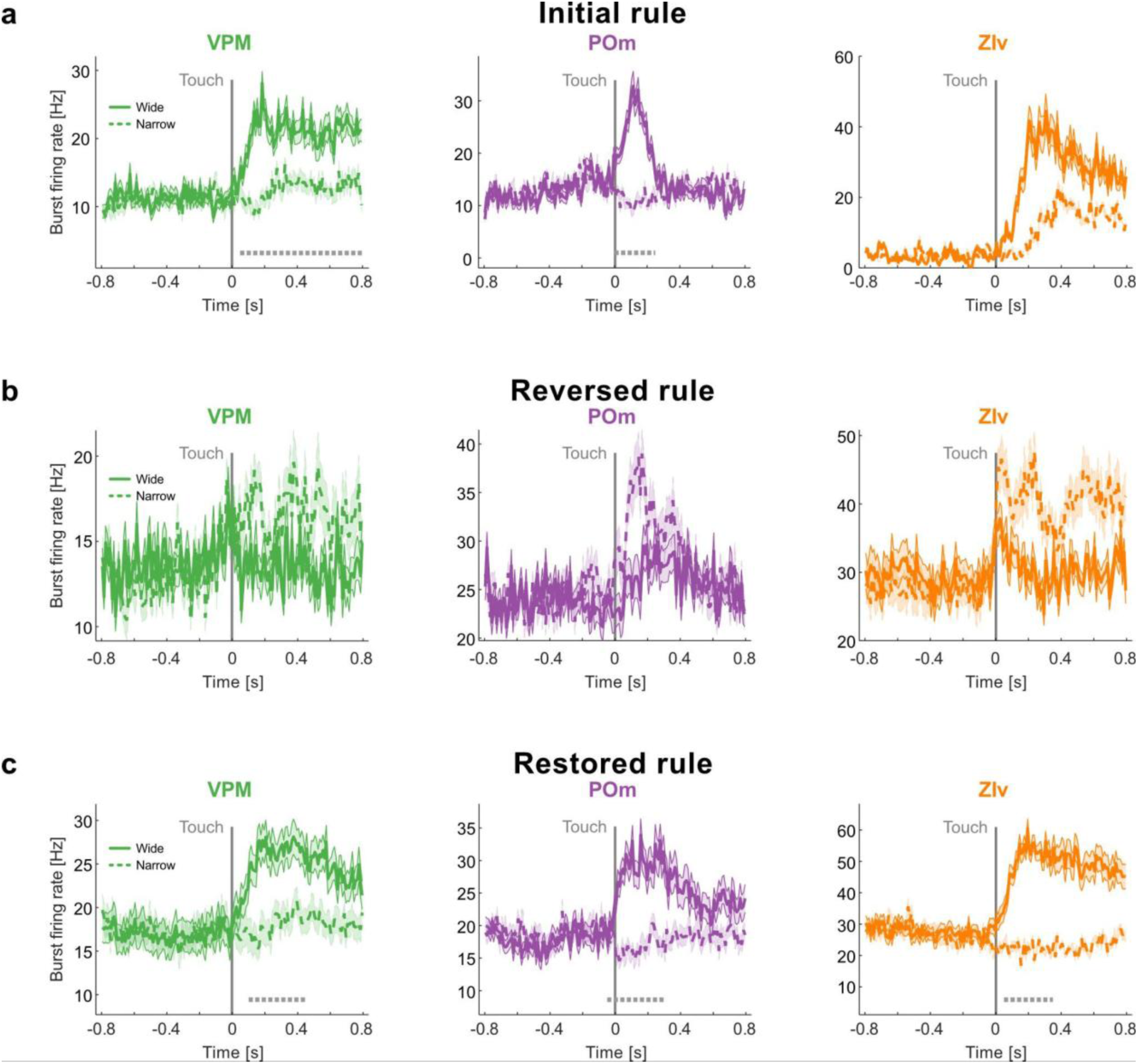
Burst firing rates during different rule stages across brain regions. Burst firing rates in VPM (left, green), POm (middle, purple), and ZIv (right, orange) in response to wide (solid line) and narrow (dashed line) apertures, represented as mean ± SEM. Statistically significant differences (α = 0.05, two-tailed Welch’s t-test) are indicated with dashed lines beneath the plot. Firing rates are shown for the initial rule (**a**), reversed rule (**b**), and the restored rule (**c**). Data pooled from n = 6 mice.

**Extended Data Fig. 5.**
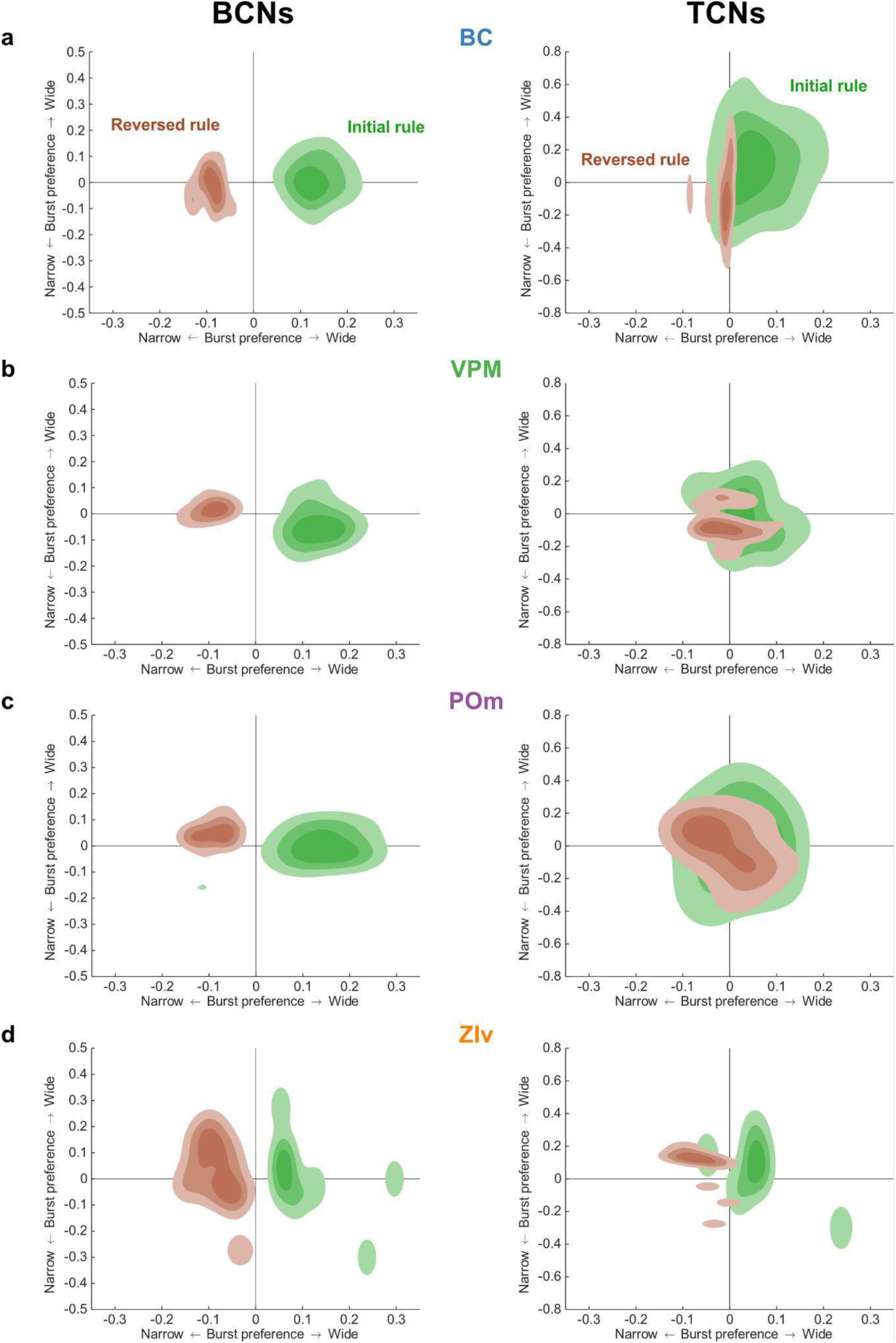
Distribution of burst and tonic bias of BCNs and TCNs across brain regions. Distributions of burst and tonic bias from BCNs (left) and TCNs (right), for BC (**a**), VPM (**b**), POm (**c**), and ZIv (**d**). The initial rule is shown in green and the reversed rule in maroon. Contour lines represent the first quartile (Q1), median (Q2), and third quartile (Q3) of the units in the scatter plot. Data pooled from n = 6 mice.

**Extended Data Fig. 6.**
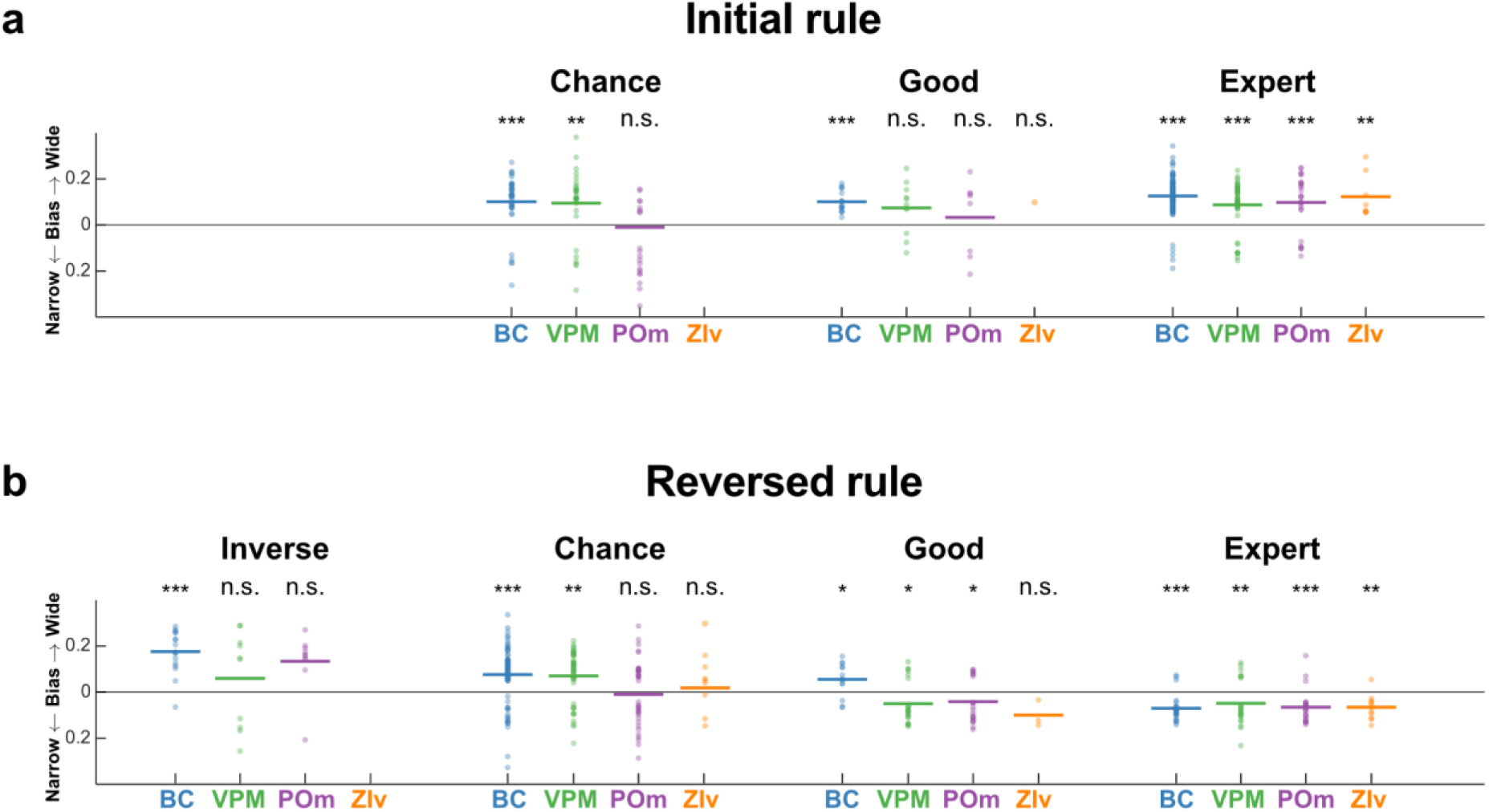
Burst bias progression for individual BCNs during learning across brain regions. Burst bias in individual BCNs of BC, VPM, POm, and ZIv. Data is shown for the learning of the initial rule (**a**) and reversed rule (**b**). The burst bias of each unit is shown across various performance categories: inverse (d’ ≤ -1.65), poor (-1.65 < d’ < 0.5), chance (-0.5 ≤ d’ ≤ 0.5), good (0.5 < d’ < 1.65), and expert (d’ ≥ 1.65). n.s.: p > 0.05, *: p ≤ 0.05, **: p ≤ 0.01 and ***: p ≤ 0.001; a,b: two-tailed Wilcoxon signed rank test. For exact p values see Extended Data Table 1. Data pooled from n= 6 mice.

**Extended Data Fig. 7.**
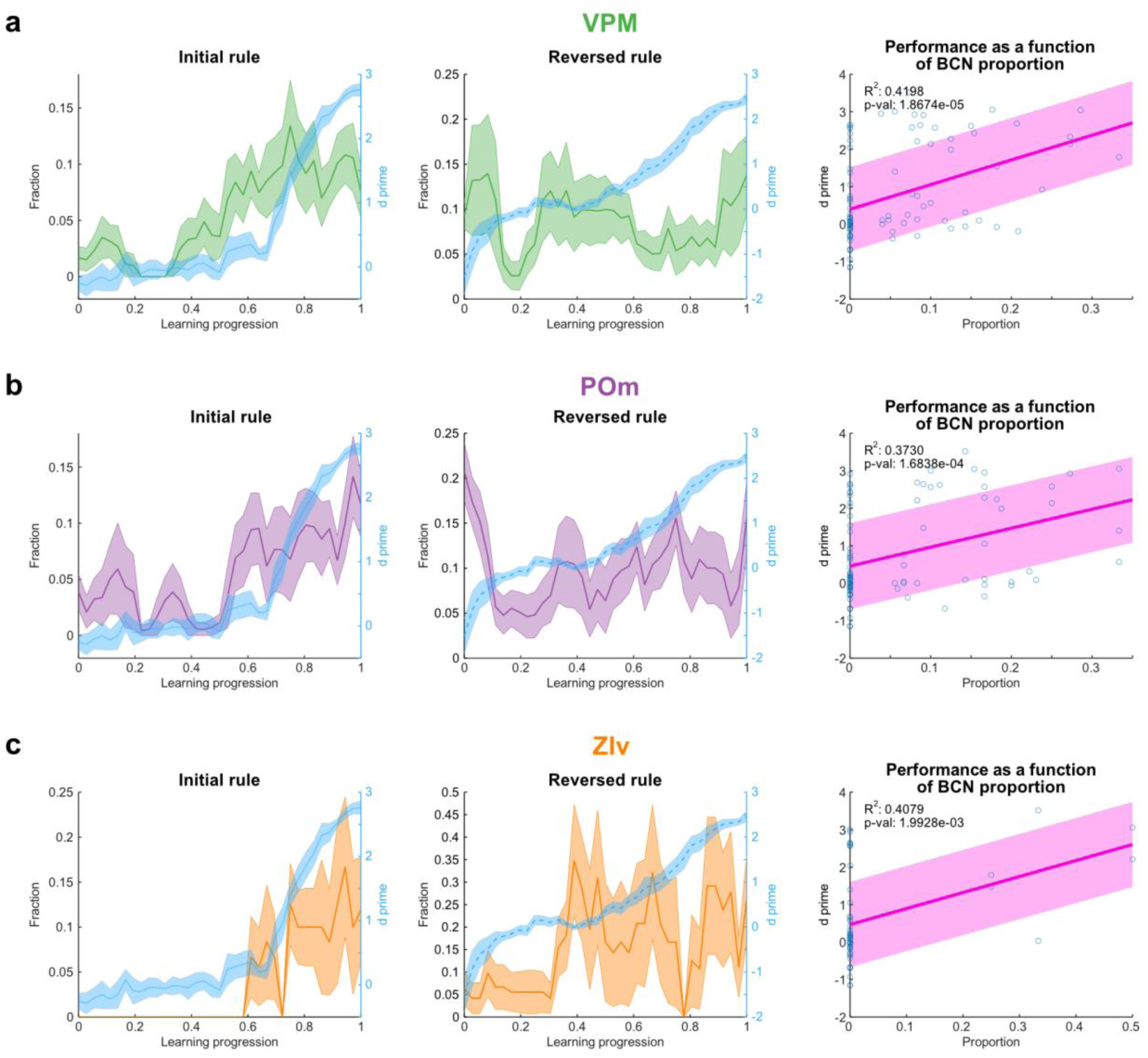
Progression of BCN proportions with improving performance across brain regions. BCN proportions during initial (left panel) and reversed rules (mid panel) for VPM (**a**), POm (**b**), and ZIv (**c**). Average BCN proportions across animals are shown as a solid line, with SEM as shaded areas. The blue dashed line represents the mean performance (d’). The right panel displays a scatter plot of session performance (d’) as a function of BCN proportion, showing a significant positive correlation in all brain regions (VPM: R^2^ = 0.4198, p = 1.8674e-05; POm: R^2^ = 0.3730, p = 1.6838e-04; ZIv: R^2^ = 0.4079, p = 1.9928e-03). Data pooled from n = 6 mice.

**Extended Data Fig. 8.**
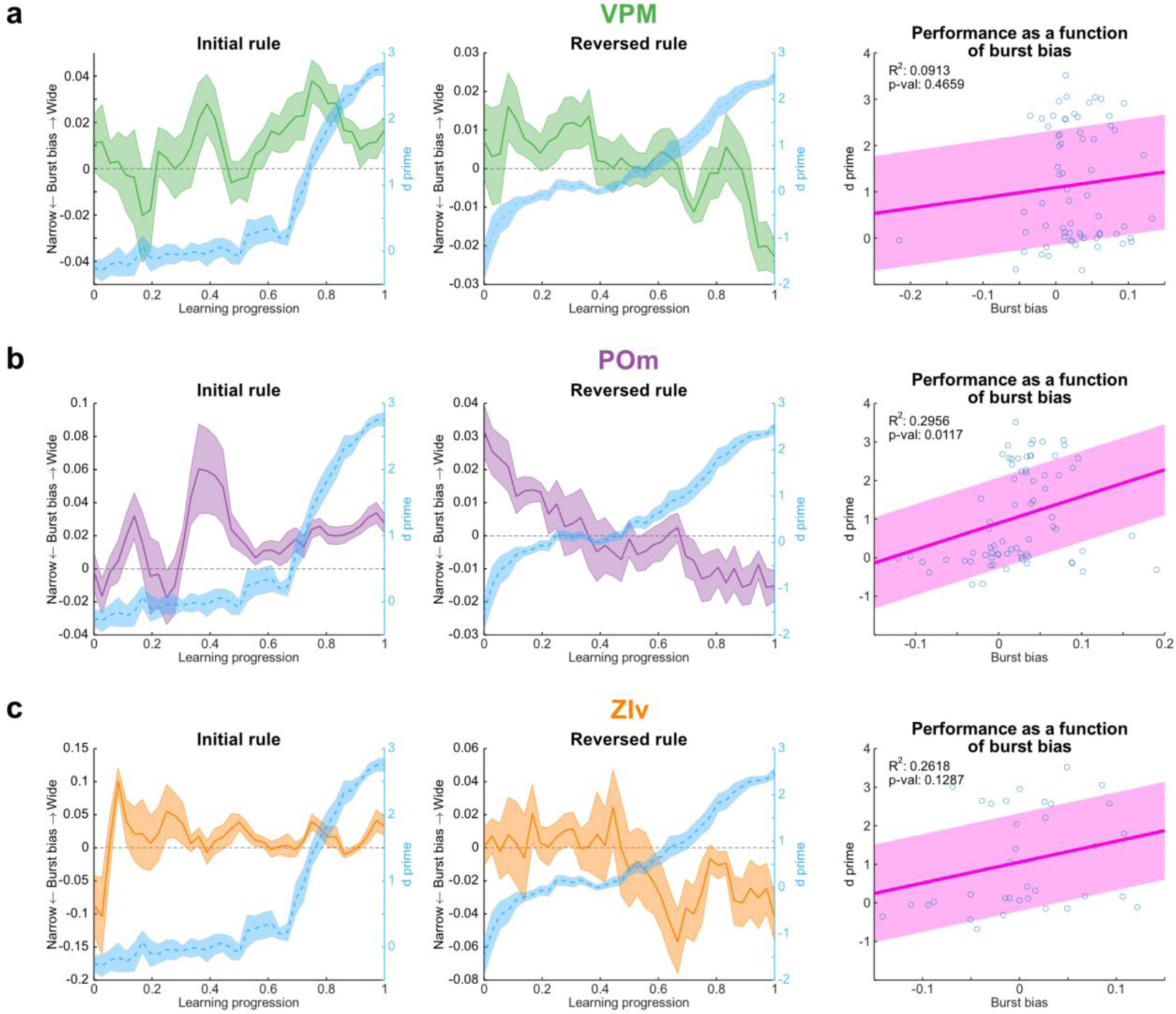
Progression of burst bias and performance across brain regions. Average burst bias of touch-modulated units during initial (left panel) and reversed rules (mid panel) for VPM (**a**), POm (**b**), and ZIv (**c**). The average burst bias across animals is shown as a solid line, with SEM as shaded areas. The blue dashed line represents the mean performance (d’). The right panel displays a scatter plot of session performance (d’) as a function of the burst bias, showing a positive correlation in all brain regions (VPM: R^2^ = 0.0913, p = 0.4659; POm: R^2^ = 0.2956, p = 0.0117; ZIv: R^2^ = 0.2618, p = 0.1287). Data pooled from n= 6 mice.

**Extended Data Supplementary Video 1:**
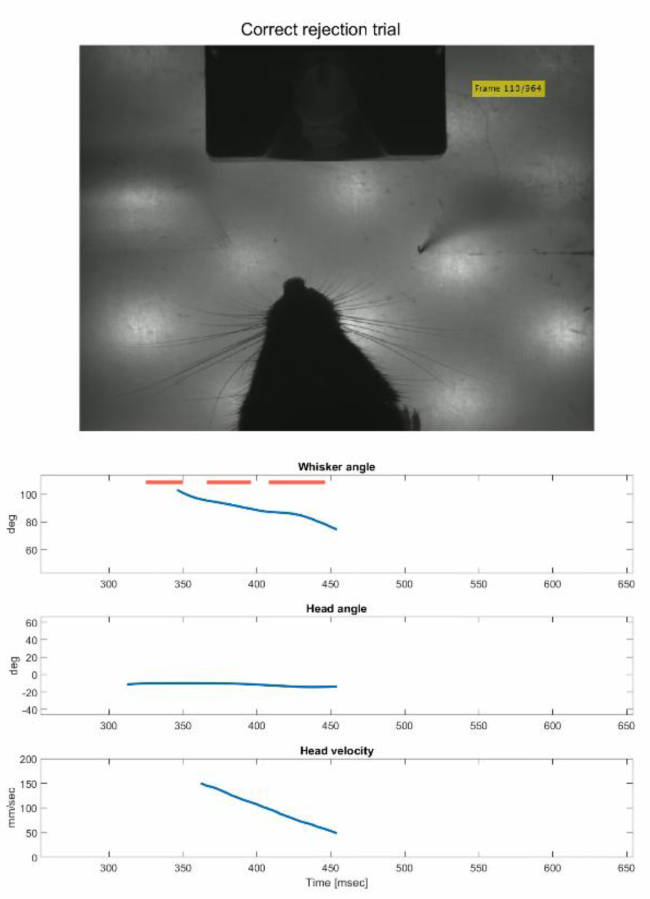
Example correct rejection (CR) trial captured with a high-speed camera (240 FPS, playback slowed by a factor of 10). The mouse successfully identifies the punished aperture and moves away from the lick port. Alongside the video, plots illustrate (1) the average whisker angle relative to the whisker pad (greater angles indicating protraction), (2) the head angle in relation to the midline (values >0° reflect a rightward tilt), and (3) the velocity of the mouse’s head within the camera’s field of view.

**Extended Data Supplementary Video 2.**
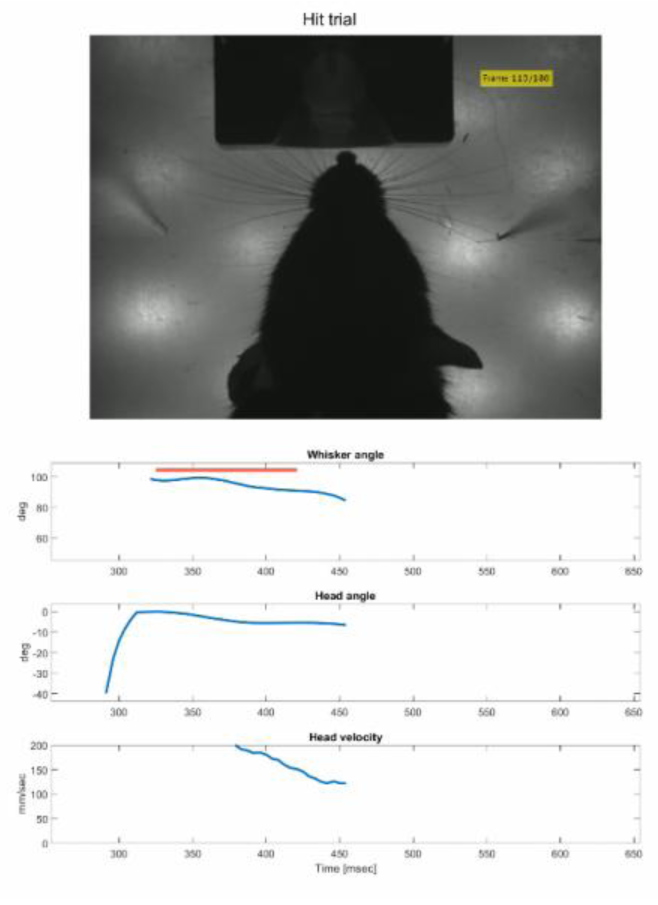
Example hit trial captured with a high-speed camera (240 FPS, playback slowed by a factor of 10). The mouse successfully identifies the rewarded aperture and licks the port to receive the reward. Alongside the video, plots illustrate (1) the average whisker angle relative to the whisker pad (greater angles indicating protraction), (2) the head angle in relation to the midline (values >0° reflect a rightward tilt), and (3) the velocity of the mouse’s head within the camera’s field of view.

**Extended Data Supplementary Video 3:**
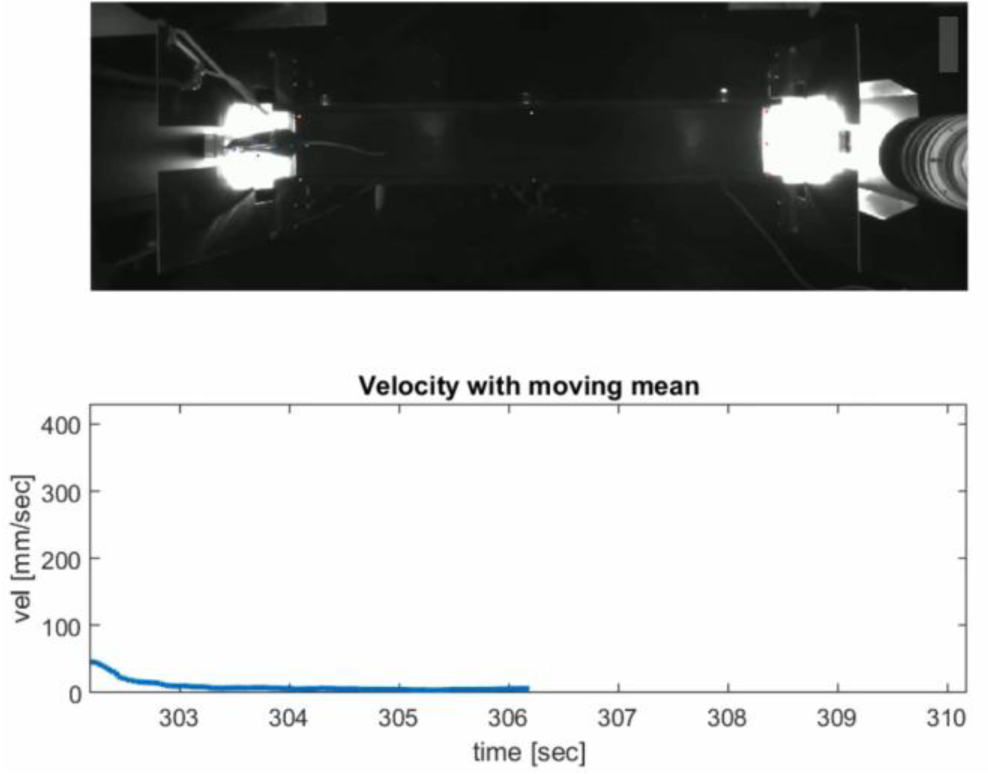
Wide-angle camera recording (60 FPS, playback accelerated by a factor of 2) providing an overview of the experimental setup. The mouse navigates a linear track, alternating between obtaining rewards and avoiding punishments based on the aperture configuration. The aperture’s state (wide or narrow) changes automatically and at random. Punishment delivery speakers are located behind the track, and high-speed cameras are positioned at the reward sites to capture whisker movements during stimulus interaction.

